# Occurrence but not intensity of mortality rises towards the climatic trailing edge of tree species ranges in European forests

**DOI:** 10.1101/2020.10.30.362087

**Authors:** Alexandre Changenet, Paloma Ruiz-Benito, Sophia Ratcliffe, Thibaut Fréjaville, Juliette Archambeau, Annabel J. Porte, Miguel A. Zavala, Jonas Dahlgren, Aleksi Lehtonen, Marta Benito Garzón

## Abstract

**Aim:** Tree mortality is increasing worldwide, leading to changes in forest composition and altering global biodiversity. Yet, due to the multi-faceted stochastic nature of tree mortality, large-scale spatial patterns of mortality across species ranges and their underlying drivers remain difficult to understand. Our main goal is to describe the geographical patterns and drivers of the occurrence and intensity of tree mortality in Europe. We hypothesize that the occurrence of mortality represents background mortality and is higher in the margin than the core populations, whereas the intensity of mortality could have a more even distribution according to the spatial and temporal stochasticity of die-off events.

**Location:** Europe (Spain, France, Germany, Belgium, Sweden and Finland)

**Time period:** 1981 to 2014.

**Major taxa studied:** More than 1.5 million trees belonging to 20 major forest tree species

**Methods:** We develop hurdle models to tease apart the occurrence and intensity of tree mortality in National Forest Inventory plots at range-wide scale. The occurrence of mortality indicates that at least one tree has died in the plot and the intensity of mortality refers to the number of trees dead per plot.

**Results:** The highest mortality occurrence was found in peripheral regions and the climatic trailing edge linked with drought, whereas the intensity of mortality was driven by competition, drought and high temperatures and was uniformly scattered across species ranges.

**Main conclusions:** Our findings provide a new perspective in our understanding of tree mortality across species ranges. We show that tree background mortality but not die-off is generally higher in the trailing edge populations, but whether other demographic traits such as growth, reproduction and regeneration would also decrease at the trailing edge of European tree populations needs to be explored.

## 1 INTRODUCTION

Tree mortality is occurring worldwide (Allen, Breshears, & McDowell, 2015; IPCC, 2014; McDowell et al., 2018). Tree mortality can change forest community, ecosystem dynamics and function, and hence alter biodiversity (McDowell et al., 2008). Yet, tree mortality remains difficult to predict at large spatial scales (Hartmann et al., 2018) because it is a multi-faceted, stochastic process (Franklin, Shugart, & Harmon, 1987). Background tree mortality is a phenomenon that generally occurs in individual trees (Hartmann et al., 2018) and is defined as the local mortality rate in the absence of catastrophic events (Csilléry, Seignobosc, Lafond, Kunstler, & Courbaud, 2013; Franklin et al., 1987; McDowell et al., 2018). It is a complex process driven by the combination of climate, forest composition, trees interactions and age (Cailleret et al., 2017; Hülsmann, Bugmann, & Brang, 2017; Ruiz-Benito, Lines, Gómez-Aparicio, Zavala, & Coomes, 2013). In contrast, die-off mortality is a local phenomenon where many trees die together in the same environment (Bugmann et al., 2019; Mueller-Dombois, 1987). Die-off mortality has been related to heatwaves and climate warming including extreme localized events, disturbances or environmental conditions such as intense drought, storms or fire (Allen et al., 2010; Breshears et al., 2005; McDowell, 2008), and is exacerbated by pest and disease outbreaks (Anderegg et al., 2015; Kurz et al., 2008).

Climate change, especially increases in the number and duration of drought events, has been linked to increases in both background mortality rates and the extent of die-off events (Allen et al., 2015; Allen et al., 2010; Benito Garzón, Ruiz-Benito, & Zavala, 2013). However, identifying the drivers of dieoff and background mortality along large environmental gradients remains challenging because tree sensitivity to biotic and abiotic factors depends on the species identity (Ruiz-Benito et al., 2013), their age (Hülsmann et al., 2017) and their ecological strategies (Benito Garzón et al., 2018; Ruiz-Benito et al., 2017; Archambeau et al., 2020).

In the context of climate change, we could expect an increase of mortality in trailing edge populations due to increased drought and rising temperatures (Benito Garzón et al., 2013; Carnicer et al., 2011; Purves, 2009; Young et al., 2017). However, very little is known about the differential drivers of background tree mortality and die-off events at large geographical scales, and both processes can occur throughout the species ranges, in the leading edge, the trailing edge and the core of the species ranges (e.g. Jump, Mátyás, & Peñuelas, 2009; Allen et al. 2010; Greenwood et al. 2017). Although, we could expect higher background mortality at the margins than at the core of the distribution (Neumann, Mues, Moreno, Hasenauer, & Seidl, 2017), and most intense events of mortality evenly distributed across species ranges (Allen et al., 2015; Allen et al., 2010; Jump et al., 2009; Jump et al., 2017).

Here, we analyze tree mortality of 20 major forest tree species from more than 1.5 million trees recorded in the National Forest Inventories from Spain, France, Germany, Belgium (Wallonia), Sweden and Finland to understand their patterns along species distribution ranges. We assume that the occurrence of mortality found in a plot reflects background mortality whereas the intensity of tree mortality found in a plot reflects die-off events. We develop hurdle models of mortality occurrence and intensity to understand the effect of climatic marginality defined as areas exhibiting the highest or the lowest values of several climatic variables and its interaction with drought. The aims of our study are to i) identify the underlying drivers of mortality occurrence and intensity and how they are influenced by the marginality of the population, and ii) to evaluate tree mortality occurrence and intensity patterns across species distribution ranges. We hypothesize that marginal populations will have higher occurrence of mortality than core populations, and that the intensity of mortality will show a patchy distribution over spatial range reflecting the stochastic nature of die-off events.

## 2 MATERIAL & METHODS

### 2.1 National Forest Inventory harmonization

We used mortality records and stand variables from National Forest Inventories (NFIs) of five countries (Spain, Germany, Finland, Sweden, Wallonia (Belgium)) (harmonized in FunDivEUROPE; (Baeten et al., 2013)) and the French National Forest Inventory (harmonized in Archambeau et al., 2020). The French NFI has temporary plots recorded between 2005 and 2014 whereas the other countries have permanent plots sampled several years apart, ranging from 1981 to 2011 (Supporting Information Table S1). Data from the six NFIs together cover a latitudinal gradient from 36° N (Spain) to 70.05° N (Finland).

### 2.2 Plot-level tree mortality recorded from NFI

We used individual tree mortality data for 20 major forest tree species, gathering a total of 1,649,850 trees and 235,394 plots (Supporting Information Table S2) varying from 10 to 263 cm (mean = 28) diameter at breast height (DBH; cm) and with a mean census intervals of 10.7 years ranging from 2 (29 plots) to 20 years (46 plots). Mortality occurrence was calculated as a binary variable, with zero when all trees were alive in the plot and one when at least one tree died in the plot during the census interval. Mortality intensity was calculated in each plot as the percentage of trees that died between the first and second inventory in the NFIs with permanent plots, divided by the number of years between census and calculated at the hectare level. In the French NFI tree mortality per plot was calculated as the percentage of trees that died within the five years before sampling in the temporary plot. We removed plots with trees recently recorded as harvested or managed between consecutive inventories and individual trees under 100 mm DBH to make the tree measurements consistent across country with different DBH thresholds (Table S1).

To avoid the potential bias caused by the different years of NFIs campaigns and the different size of the plots between the NFIs (Table S1), we upscaled tree mortality from plot to hectare per year using a weighted index provided by each NFI and dividing this value by the number of years between campaigns for NFI with repetitive measurements or by five for the French NFI, where mortality was estimated for five years (Supporting Information Table S3). The weighted index reflected the size of the plot or the density of the grid or both depending on the country (Table S1 and http://project.fundiveurope.eu/).

### 2.3 Model predictors

#### 2.3.1 Indexes of climatic marginality and climatic areas

We determined the distribution range of each species using information available from Caudullo, Welk, & San-Miguel-Ayanz (2017) or EUFORGEN (http://www.euforgen.org/). Within each range we characterised the climate using a Weighted Principal Component Analysis (WPCA, Benito-Garzón, Leadley, & Fernández-Manjarrés, 2014) based on 21 climatic variables averaged over the 2000-2014 time period at each point in a 1 x 1 km pixel size grid (Supporting Information Table S4). The WPCA was calculated using 10,000 randomly selected points within each species ranges. The variance explained by the two first axis of the WPCA ranged from 71.53% to 87.42% for *Fagus sylvatica* and *Larix decidua*, respectively, Supporting Information Table S8). Based on the weighted scores of the two first WPCA axis we defined three climatic groups: core, transition and marginal regions (Supporting Information Figure S1 and Table S5). Species-specific thresholds for attributing the core (C), climatic marginal (M) and transition (T) areas were calculated based on the WPCA scores (Table S2): values from 0 to 60% were attributed to core areas, values between 60 and 80 % are transition areas and values higher than 80% are marginal areas.

To further separate climatic marginal areas (M) into climatic trailing edge (TE) for the southernmost one and climatic leading edge (LE) for the northernmost, we used a Discriminant Principal Component Analysis (DPCA) and an attribution test to check whether individual points were successfully reassigned in their attributed group based on the discriminant functions (Jombart, 2008; Figure S1).

Finally, NFI plots were linked with the WPCA scores and classified as core (C), leading or trailing edge (LE and TE) accordingly. Plots lying in the transition region (T) were not used in the analysis (Supporting Information Table S2 and Figures S1,S2).

#### 2.3.2 Climatic data

We characterised the long-term climate of each plot with eight climate variables. To make the variables comparable between different survey dates and countries, we averaged them over the last 30 years prior to the first survey (hereafter climatic variables; Table S4) (Fréjaville & Benito Garzón, 2018). The eight variables included four temperature-related variables and four precipitation-related variables that were uncorrelated and that have been shown to have an effect on tree mortality (Archambeau et al., 2020; Benito Garzón et al., 2018; Ruiz-Benito et al., 2017): annual mean temperature (bio1), maximal temperature of the warmest month (bio5), winter mean temperature (tmean.djf), autumn mean temperature (tmean.son) (temperature variables) & annual precipitation (bio12), precipitation of the wettest month (bio13), precipitation of the driest month (bio14), annual water balance (precipitation minus potential evapotranspiration (ppet.mean) (Table S4).

In addition, we used the Standardized Precipitation Evapotranspiration Index (SPEI v.2.5 (2017) (http://hdl.handle.net/10261/104742). SPEI is a multi-scalar drought index where negative values indicate drier conditions over the timescale considered (from 3 to 48 months), relative to median values for a long-term reference period (from 1901 to 2015) (Vicente-Serrano, Beguería, & López-Moreno, 2010). For each month during the time interval between inventory campaigns, we used 1901 to 2015 as a reference period and calculated SPEI monthly index considering a period of 12 months relative to our reference period. For each plot and based on these monthly data, we calculated the annual means and extracted the minimum and mean values for this time period (hereafter SPEI variables; Table S4).

#### 2.3.3 Stand and competition variables

All stand variables were calculated using NFI data, transformed where necessary to meet the model assumptions of normality (Supporting Information Table S6): total basal area increment (BAI_j_, m^2^ ha yr^−1^), calculated as the difference in basal area between two inventory periods for all NFIs except France where five years cores were used; mean basal area increment (meanBAI_j_, m^2^ ha yr^−1^); mean diameter at breast height (DBH, mm), tree density calculated as the number of trees per hectare (treenumber, No. trees ha^−1^); total (BA, m^2^ ha^−1)^ conspecific stand basal area (estimated as the basal area of all individuals of the species in the plot, BAcon, m^2^ ha^−1^) and heterospecific stand basal area (estimated as basal area of all individuals excluding the studied species) (BAhetero, m^2^ ha^−1^).

The DBH, BAI_j_, meanBAI_j_ and treenumber were included in the model as proxies of the average age (DBH), growth and tree density in the plot because they are known to influence tree mortality (Hülsmann et al., 2017; Vanoni, Cailleret, Hülsmann, Bugmann, & Bigler, 2019). The number of years between surveys (yearsbetweensurvey) was also included in the model to account for mortality probability increases with elapsed time. We used BA, BAcon and BAhetero as proxies of total competition, intraspecific and interspecific competition (Kunstler et al., 2016).

### 2.4 Statistical analyses

#### 2.4.1 Selection of climatic and competition covariates in the mortality models

For each species, we ran 48 competing occurrence of mortality models. In each model we included the climatic marginality as a qualitative variable (i.e. the core, leading or trailing edge of each plot), the five stand covariates, and the minimum and mean SPEI indexes. We added all the possible combinations of one precipitation-related, one temperature-related and two competition variables. We included all interactions between marginality, the two SPEI indexes, the two competition-related variable and the two climate variables (Table S7).

We included both precipitation- and temperature-related variables to the models, in addition to marginality, because they could vary within the species margins and thus capture variations not accounted by marginality variable. The collinearity between precipitation- and temperature-related variables and marginality were assessed using variation inflation factors (Supporting Information Table S8a,b).

#### 2.4.2 Statistical models of mortality

We used species-specific hurdle models to handle the zero-inflated distribution of tree mortality (Ruiz-Benito et al. 2017; Benito Garzón et al. 2018; Archambeau et al. 2020). Consequently, we analyzed separately the mortality occurrence between two census (0/1 = at least one tree is dead in the plot/ all trees are alive in the plot) and the intensity of mortality in plots where mortality occurs (the proportion of trees dead in the plot, Young et al., 2017). Firstly, mortality occurrence was analyzed with the binomial part of the hurdle model (*Y* 1*_i_*=1, table S3) where *p_i_* is the probability of occurrence of a mortality event in an individual plot *i* during the census interval. We used a binomial GLMM with a *logit* link (BIN model) to estimate the parameters of the species-specific linear function *η*_1 *i, sp*_ (Hülsmann et al., 2017):

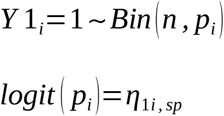

Secondly, we analyzed the intensity of mortality as the annual rate of mortality in plots where at least one tree was recorded as dead (*Y* 2_*i*_, Table S3) with zero truncated negative binomial mixed-effect models in the second part of the hurdle model (NB model), *Y* 2_*i*_ where *μ_i_* is the mean number of mortality events per year per hectare and *k* is the inverse of the dispersion (*Y*2*_i_*~*NB*(*μ_i_*, *k*)). We used NB models with a *log* link to estimate the parameters of the species-specific linear function *η*_2*i, sp*_:

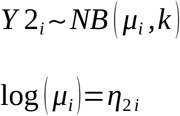

Functions *η*_1 *i, sp*_ and *η*_2*i, sp*_ take the same general form:

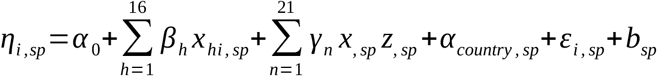

Where *α*_0_ is an intercept term, *α_country, sp_* is the random country intercept to account for sampling differences between each NFI and follows a Gaussian distribution 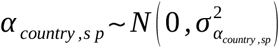; *ε_i, sp_* is the residual error following a Gaussian distribution 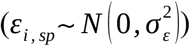; *b_sp_* is an autocorrelation spatial effect that follows a Matérn distribution (*b_sp_~Matérn*(*v_sp_, ρ_sp_*), where *v_sp_* is the smoothness and *ρ_sp_* the shape, Supporting Information Figure S3). *β_h_* is the regression coefficient for the *h*th of 16 fixed effect predictors *x_sp_* (including 5 stand covariates, 2 climatic variables and their respective quadratic effect, 2 drought-related (SPEI) variables and their respective quadratic effect, 2 competition variables and marginality, see details below and in Supporting Information Tables S4, S6 and S7) and *γ_n_* the regression coefficient of the *n*^th^ interaction between fixed effect predictors *x_sp_* and *z_sp_* (including all interactions between climatic variables, drought-related variables, competition variables and marginality).

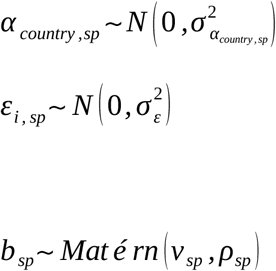

#### 2.4.5 Model selection

To select the most parsimonious model, we applied the following procedure for each species: (1) We calculated the Variance Inflation Factor (VIF) for all 48 possible combinations of variables and removed combinations with VIF > 10 (Dormann et al., 2013; Table S8a,b); (2) We ran BIN models including each remaining combinations of variables and selected the combination with the best predictive ability using the AIC (AIC < 2) and the log H-likelihood (largest values;Lee & Nelder, 2018); (3) We fitted an NB model including the same variables as those in the BIN model with the best predictive ability. (4) We used a stepwise approach for both the BIN and NB models (i.e. we removed the least significant variable to fit a new model) to obtain the most parsimonious models. All models were fitted with the SpaMM package (Rousset, Ferdy & Courtiol, 2016; Table S6) under the R version 3.6.1.

#### 2.4.6 Model validation

The goodness-of-fit was evaluated with the area under the curve (AUC) for BIN models (Hurst, Allen, Coomes, & Duncan, 2011) and with cross-validation for the NB models (models were fitted on 66% of the data while the remaining 33% were used to validate the predictions, Table 1).

**Table 1:**
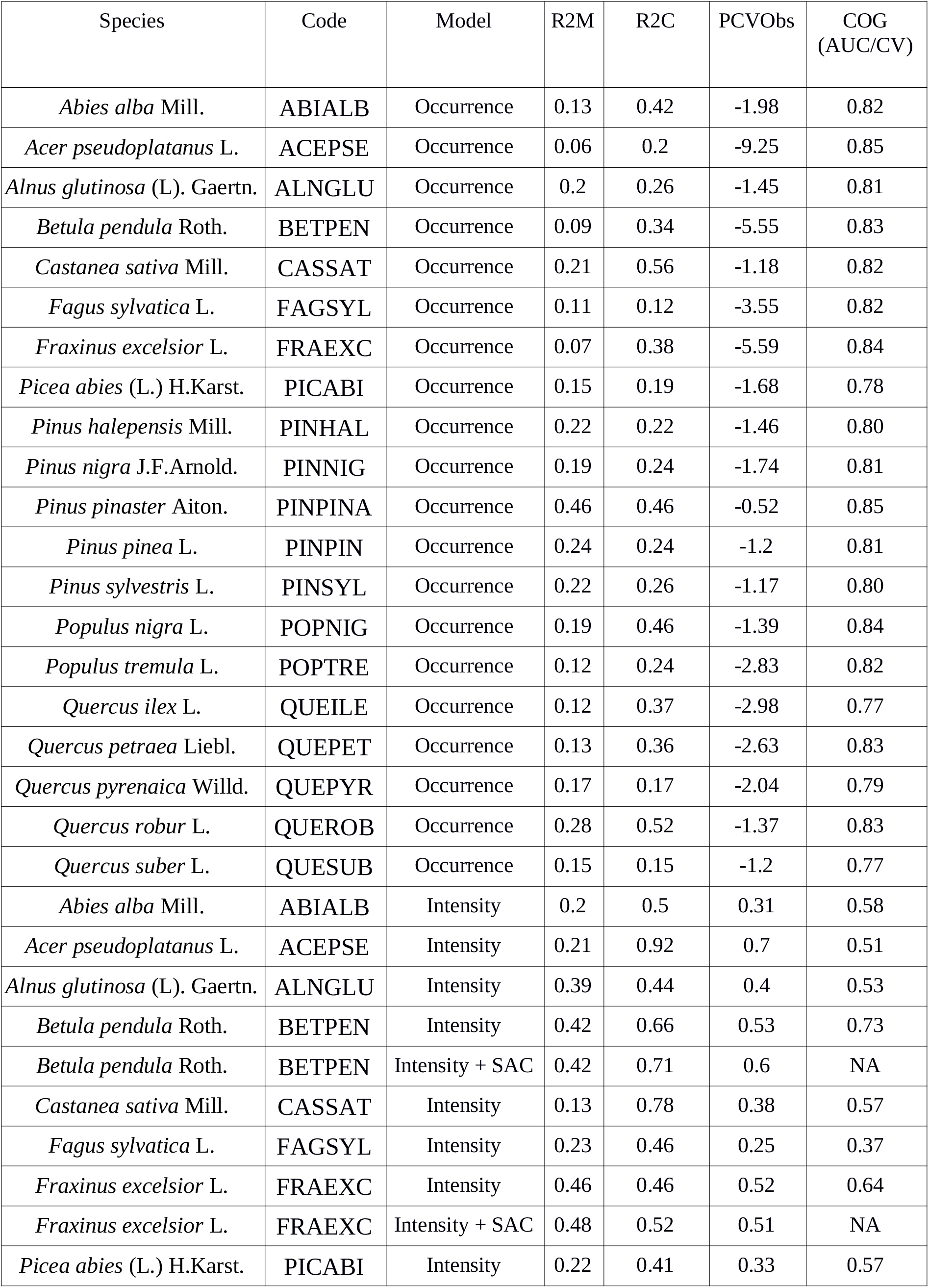
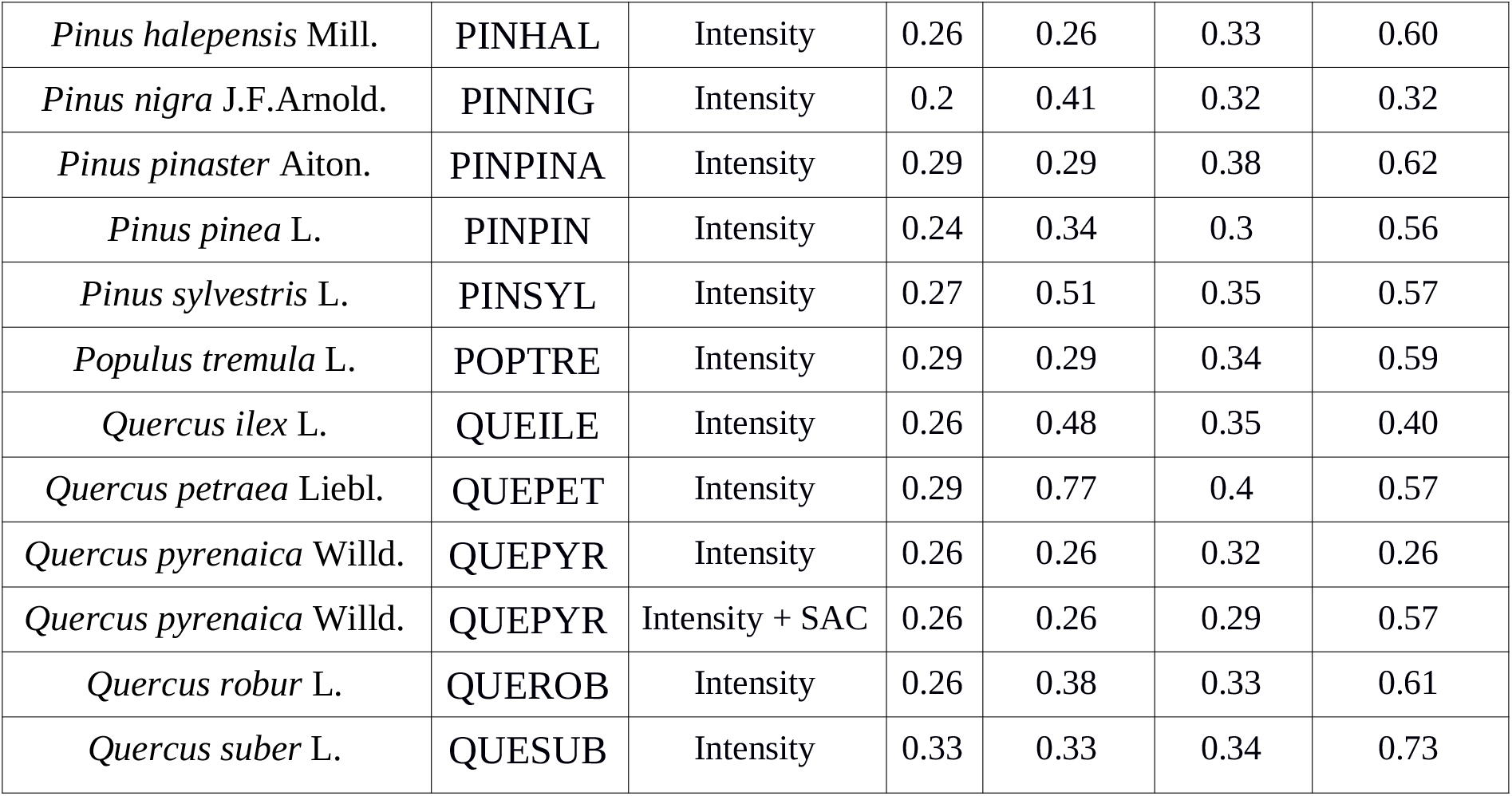
Statistical evaluation of occurrence and intensity of mortality models, by species. Species: Name of the species. Code: Code used for each species Model: model type: Occurrence, intensity or intensity of mortality model including spatial autocorrelation (intensity + SAC - Spatial Autocorrelation -). R2M: Marginal r-squared. R2C: conditional r-squared. PCVObs: proportion change in variance between null model and fixed effect model (in %). COG: capacity of generalization. AUC (Area Under the Curve) for mortality occurrence models and CV (Cross Validation score) for intensity of mortality models.

The percentage of the variance explained by BIN and NB models was estimated by the marginal and conditional R-squared including fixed-effect and fixed plus random effects, respectively (Nakagawa & Schielzeth, 2013). The proportion of change in explained variance between full and null model (PCV) indicates the variance retained by the selected model. All these metrics were calculated from the SPAMM objects using a personal script adapted from the piecewise Package (Lefcheck, 2016) according to Nakagawa methodology (Nakagawa & Schielzeth, 2013, Nakagawa, Johnson & Schielzeth, 2017).

#### 2.4.7 Comparison of spatial predictions and climatic marginality

We used the selected models to predict mortality occurrence and intensity across the range of NFI plots. To visually inspect the differences in the climatically marginal populations, we split the predicted values into three groups based on the quartiles, to indicate high, medium and low levels of mortality (Figure 3 and Supporting Information Tables S9, S10).

To statistically test for heterogeneity in the distribution of the predicted probability among the three areas (core, leading edge and trailing edge), we compared the predicted distribution (Figure 3) against the expected distribution under the assumption of no spatial structure in mortality occurrence (null hypothesis) with a χ-square test. Under the null hypothesis, we expected the distribution to be uniformly distributed within the three areas (25% of the values in each quartile) (Supporting Information Figure S4a). P-values < 0.05 indicate that predicted mortality was different than expected under the null hypothesis. The same approach was used to test for patterns across the three areas in predicted mortality intensity (Supporting Information Figure S4b).

## 3 RESULTS

### 3.1 Climatic marginality across species ranges

The variables that contributed the most in defining the core, trailing and leading areas were the annual evapotranspiration for 10 out of the 20 species, the maximum temperature of the warmest month for four species and annual precipitation for four species (Table S5). Marginal areas (leading and trailing edges) were therefore areas exhibiting the highest or the lowest values for these climatic variables.

We observed that our climatic marginality did not systematically match with the commonly used geographic marginality (northern part of species distribution corresponding to the geographical leading edge, and southern part to the geographical trailing edge), particularly in the mountainous areas which were included in the climatic leading edge for most species although they were mostly located in the central part of the range (geographical core) (Figure S2).

### 3.2 Underlying drivers of occurrence and intensity of tree mortality

Overall, the occurrence of mortality was mostly located at the trailing edge and related to drought whereas the intensity of mortality was related to multiple drivers at the trailing edge and to the warmest temperature at the leading edge. Bigger trees (likely to be older trees) showed higher occurrence of mortality than smaller ones whereas intensity of mortality was similar across sizes (and therefore ages).

The variance explained by BIN models ranged from 6% for *Acer pseudoacacia* to 46% for *Pinus pinaster* and the AUC ranged from 0.769 for *Quercus ilex* to 0.850 for *Acer pseudoplatanus* (Table 1). The variance explained by NB models ranged from 13% for *Castanea sativa* to 48% for *Fraxinus excelsior* and the cross-validation scores ranged from 0.256 for *Quercus pyrenaica* to 0.735 for *Betula pendula*. Accounting for spatial autocorrelation (SAC) in the models improved the capacity of generalization of the three models (i.e. from 25.61% to 57.54% for *Quercus pyrenaica*, Table 1).

Competition-related variables (stand basal area of the species and stand basal areas of the other species at the plot) were the most frequently significant variables in BIN and NB models (significant in 13 out of 20 species for interspecific competition and 10 species for intraspecific competition, see estimated coefficients and their standard error in Supporting Information Table S11a,b and frequency in Table S12a,b). Precipitation, temperature and SPEI variables were retained in BIN models for 16, 11 and 12 species respectively and in NB models for 11, 8 and 9 species (estimated coefficients and their standard error of the associated coefficient in Table S11a,b and frequency in Table S12a,b).

Climatic marginality was significant in the BIN model in 5 species in the trailing edge and in 7 species in the leading edge; and it was also significant in NB models in 3 species in the trailing edge and in 7 species in the leading edge (magnitude and standard error of the associated coefficient are shown in Table S11a,b and frequency in Table S12a,b).

The average age of the plot (meanDBH) was significant in explaining both occurrence and intensity of mortality (positive association in 14 species and 11 species respectively) meaning that larger (and therefore older) trees were more affected by mortality. Average growth rate of the species of the plot (meanBAIj) was also significant in both models (negative associations in 18 species for BIN models and 7 in NB models) meaning that older plots (with the lowest average growth) are more likely to experience mortality. Tree density (treenumber) was positively associated in both models (14 and 12 species respectively for BIN and NB models). The number of years between surveys yearsbetweensurvey) was also significant in explaining both the mortality occurrence probability between two census (6 species, BIN model) and the intensity of mortality (15 species, NB model) (Magnitude and standard error of the associated coefficient in Table S11a,b and frequency in Table S12a,b)

Interactions between marginality and SPEI variables were the most frequent in BIN models (17 significant interactions, 9 at the trailing edge and 7 at the leading edge) whereas the interaction between climatic marginality and temperature-related variables was the most frequent in NB models (12 significant interactions, 3 at the trailing edge and 9 at the leading edge) (Supporting Information Table S11c,d). When drought conditions in the studied period were higher than in the reference period (lower values of mean SPEI), predicted tree mortality occurrence probability (BIN models) was higher in marginal than core areas, particularly at the trailing edge (for both temperate and Mediterranean species *Abies alba, Picea abies, Pinus sylvestris, Castanea sativa Pinus pinea* and *Pinus nigra*, Figure 1a-f). Under drier conditions than those experienced in the reference period (i.e. negative SPEI values), the highest probability of mortality was found in the core area for some temperate (*Populus tremula, Quercus robur, Betula pendula* (Supporting Information Figure S5a-c) and Mediterranean species (*Pinus halepensis* and *Quercus pyrenaica* (Supporting Information Figure S5d and Table S12c).

**Figure 1:**
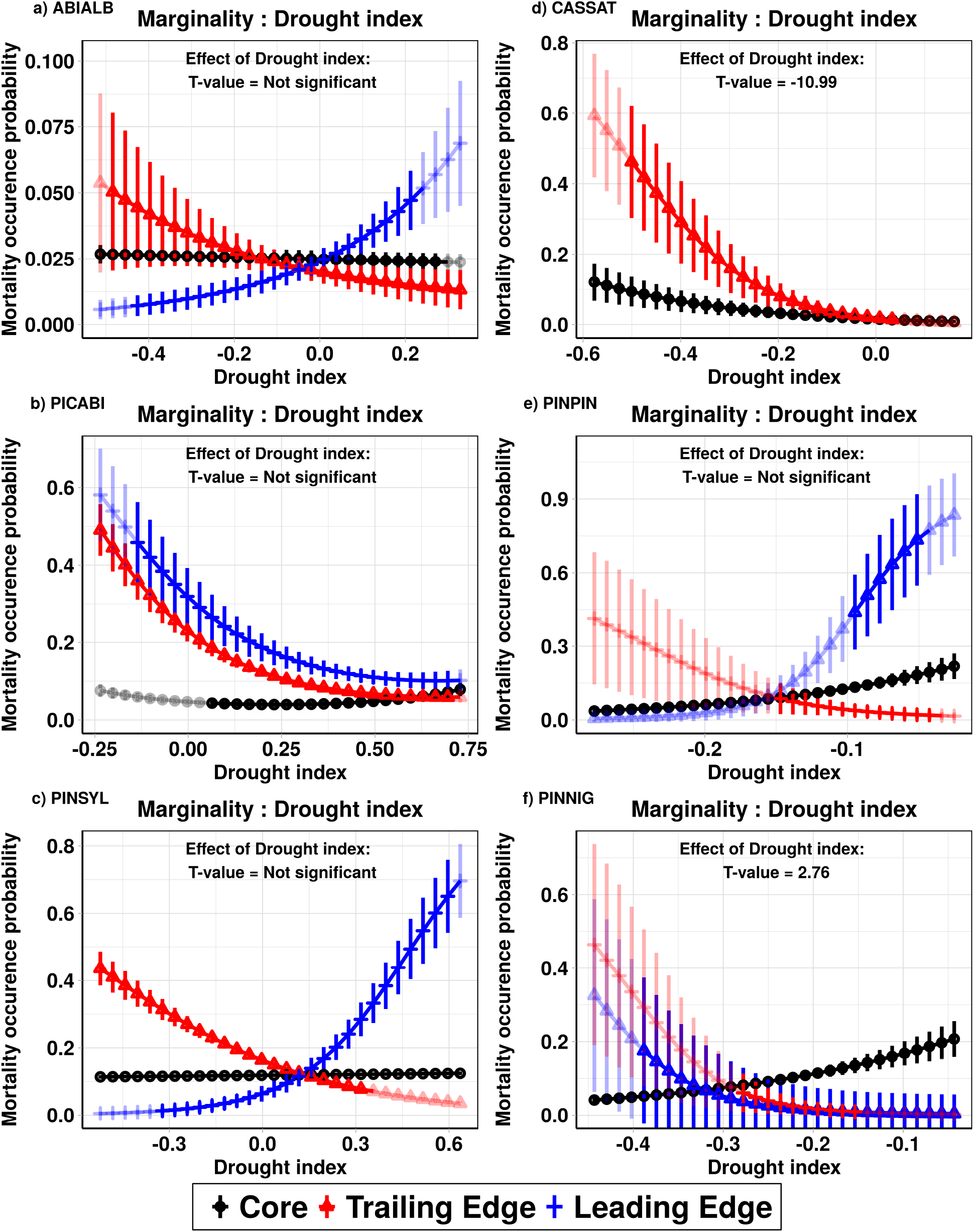
Effect of the interaction between drought (mean SPEI index) and marginality on predicted mortality occurrence per plot (expressed as a probability) across the core (black lines), trailing (red lines) and leading edge (blue lines) of three temperate species: a) *Abies alba*, b) *Picea abies*, c) *Pinus sylvestris*; and three Mediterranean – warm temperate species: d) *Castanea sativa*, e) *Pinus pinea*, f) *Pinus nigra*. Predictions within the ranges of the environmental gradients covered by the species are shown by solid colors and extrapolations outside the environmental gradients covered by the species are shown in transparent colors. For the case of *Castanea sativa* our data did not cover the leading edge of the species. T-values of the main effects interacting with marginality are reported.

In addition to drought, a few species also showed the highest tree mortality probability during the time between surveys at the climatic margins when precipitation were low or when temperatures were high. This was the case for the temperate species *Fagus sylvatica* and *Fraxinus excelsior* (Supporting Information Figure S6a and Table S12c) and the Mediterranean species *Pinus pinaster* and *Quercus suber* (Supporting Information Figure S6b-c).

The intensity of mortality (NB models) for the temperate species *A.alba, B.pendula, F.sylvatica* and *P.abies* was generally higher in the trailing edge than in the core but not always associated with low SPEI. It was associated with various variables as competition, lower SPEI, warmer temperatures or lower precipitation than in the core, depending on the species (Figure 2a-c and Supporting Information Figure S7a). In addition, a strong effect of high temperatures was observed in leading edge areas. Under high temperatures, both temperate (*F.excelsior, Quercus petraea* and *Q.robur*, Figure 2d-f) and Mediterranean species (*Quercus ilex, P.halepensis, P.nigra* and *P.pinea*; Supporting Information Figure S7b-c and Table S12d) showed more intense predicted mortality at the leading edge than in the core areas. Also, *P.tremula* populations are more likely to show high intensity of mortality at the core of its distribution when drought in the studied period was higher than in the reference period (lower values of mean SPEI) (Table S12d).

**Figure 2:**
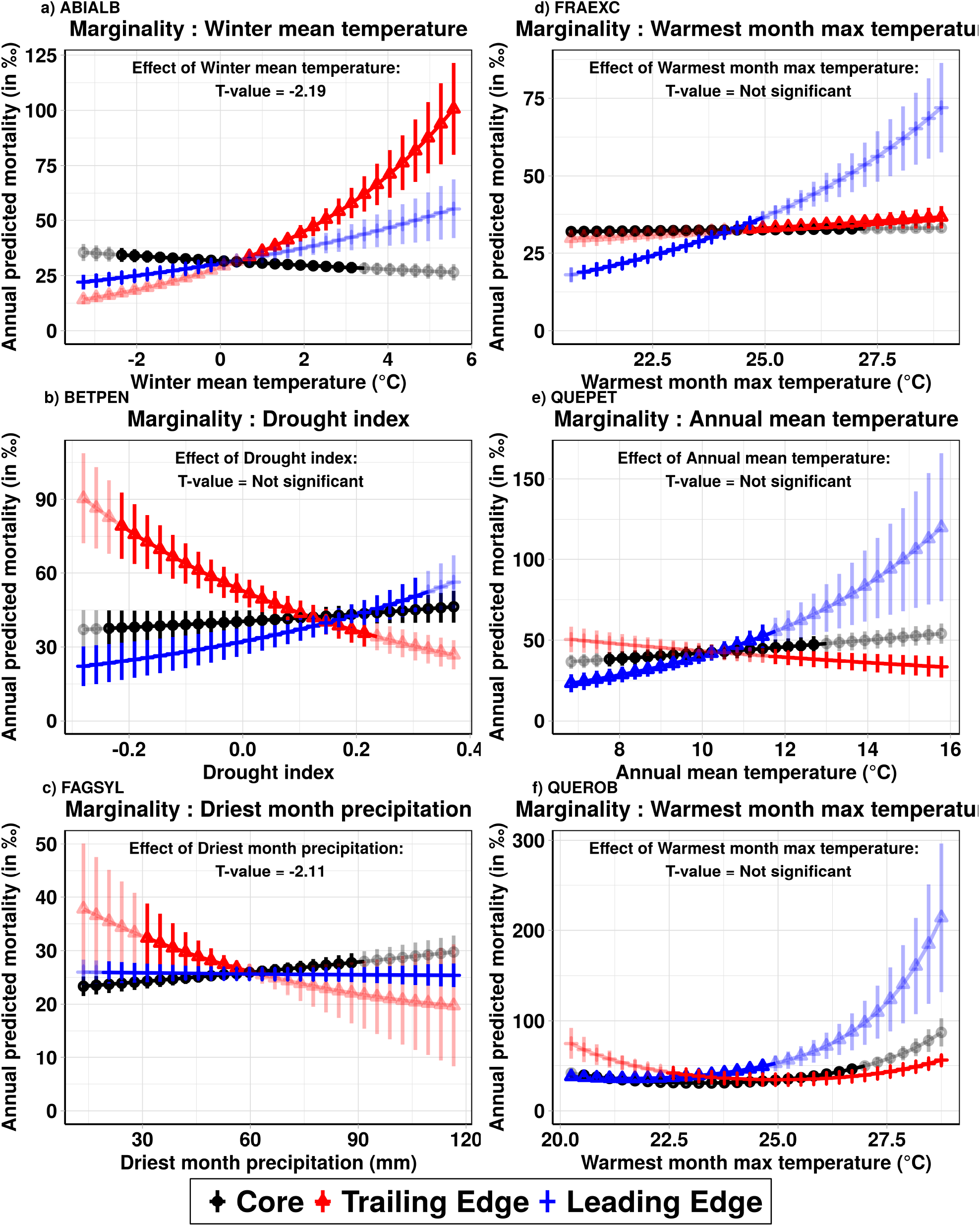
Effect of the interaction between climatic variables on the predicted intensity of mortality across the core (black lines), trailing (red lines) and leading edge (blue lines) of six temperate species: a) *Abies alba*, b) *Betula pendula*, c) *Fagus sylvatica*, d) *Fraxinus excelsior*, e) *Quercus petraea*, f) *Quercus robur*), expressed as the proportion (‰), by year and by plot. Predictions within the ranges of the environmental gradients covered by the species are shown by solid colors and extrapolations outside the environmental gradients covered by the species are shown in light colors. T-values of the main effects interacting with marginality are reported.

### 3.3 Spatial patterns of occurrence and intensity of mortality across tree species ranges

The predicted probability of mortality occurrence (BIN models) was the highest in the trailing edge part of the range for eight temperate (*Alnus glutinosa*, *Betula pendula*, *Picea abies*, *Pinus sylvestris*, *Populus tremula*, *Fagus sylvatica, Quercus robur, Q. petraea*) and three Mediterranean species (*Quercus ilex*, *Q.pyrenaica*, *Castanea sativa*) (Figure 3 and S4a). None of the species had a significantly higher probability of mortality occurrence in the core than in the margins, while two temperate species (*P.abies* and *P.sylvestris*) had lower probability of occurrence of mortality in the core than expected under the assumption of uniformity across the species range (Figure 3 and S4a). Four temperate species (*Alnus glutinosa*, *Pinus sylvetris*, *Populus nigra*, *Quercus robur*) and four Mediterranean species (*Castanea sativa*, *Pinus nigra*, *P. pinea* and *Quercus ilex*) had a lower probability of occurrence of mortality in the leading edge part of their range than the core (Figure 3 and S4a). We did not find any spatial patterns in the intensity of mortality (NB models) in temperate species. However, the highest predicted intensity of mortality was at the leading edge part of the species range for 6 Mediterranean species (*P.halepensis, P.nigra, P.pinaster, P.pinea, Q.pyreneica, Q.suber*) (Figure 3 and S4b)

**Figure 3:**
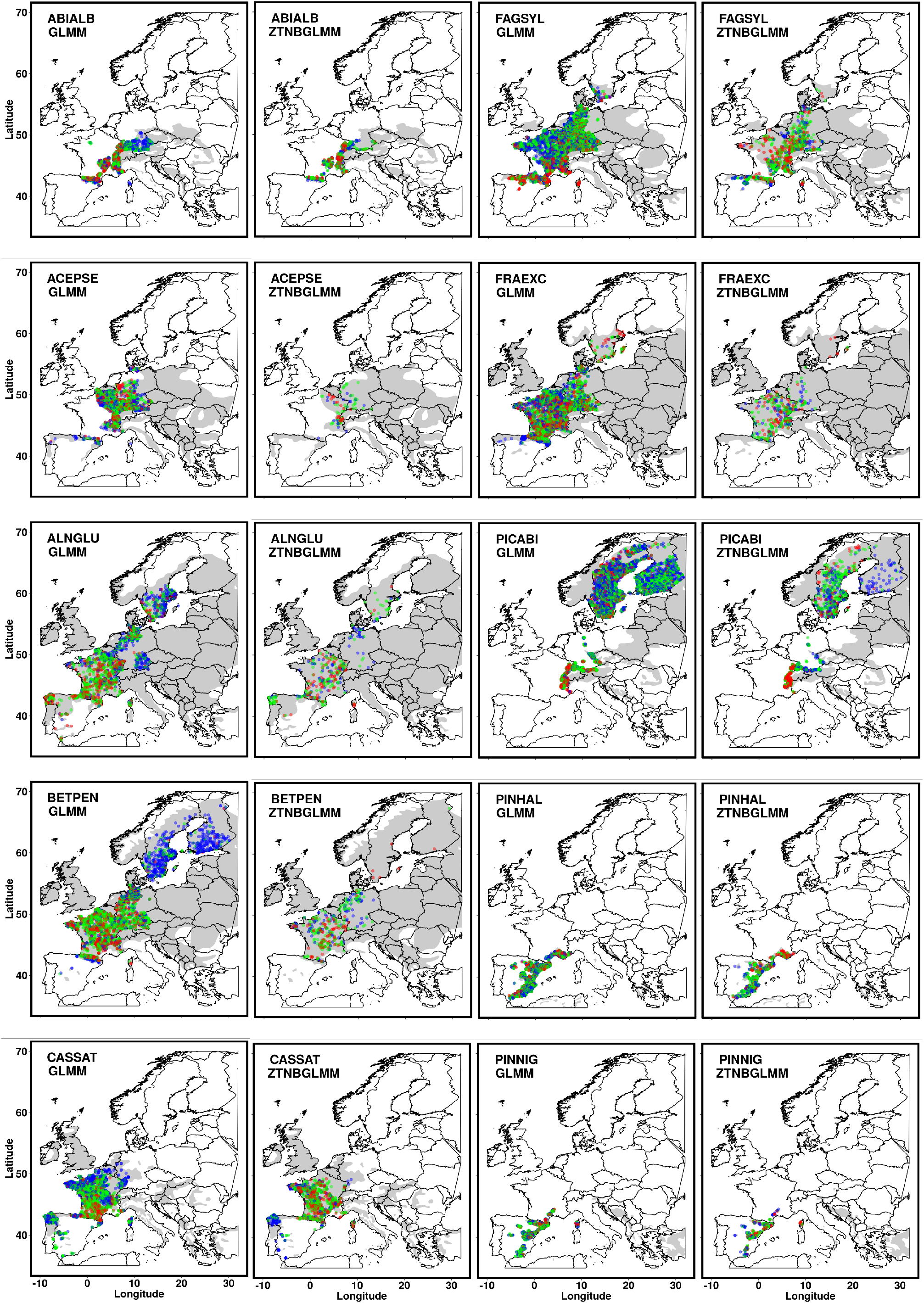
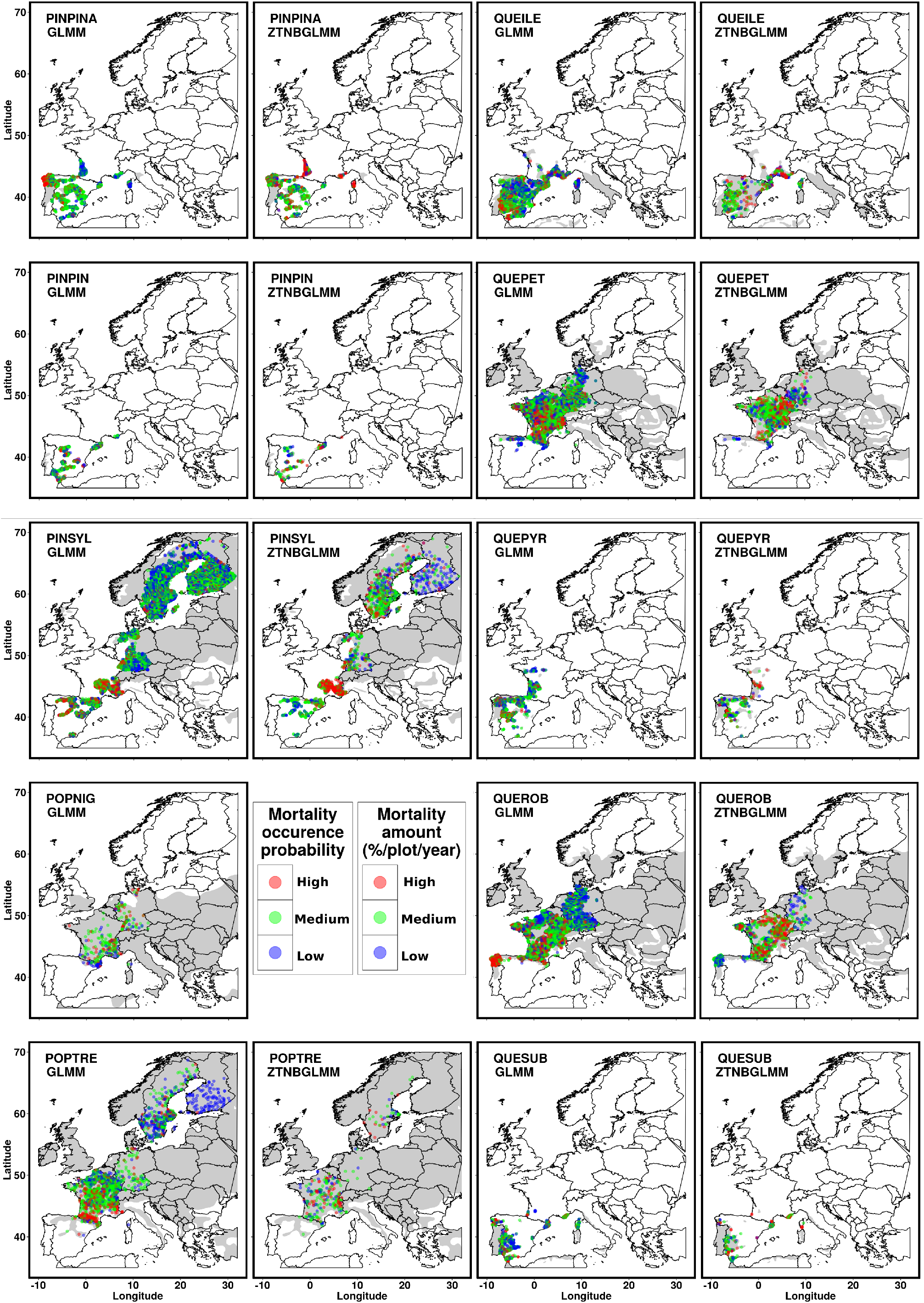
First and third columns show the predicted mortality occurrence (GLMM) and second and fourth columns are the predicted intensity of mortality (ZTNBGLMM) for the twenty species. Blues dots correspond to mortality predictions values lower than the first quartile (lowest values), green dots represent values ranging from the first to the third quartile (medium values) and red dots represent values higher than the third quartile (highest values) (Table S10). Light grey areas display species distribution ranges.

Only six species showed similarities between the occurrence and intensity of predicted mortality (Figure 3): *Castanea sativa, Quercus robur* and *Quercus ilex* showed the lowest values of both types of predicted mortality at their leading edge whereas *Acer pseudoplatanus* displayed the lowest values at the trailing edge; the predicted intensity and occurrence of mortality predictions for *Quercus ilex* showed the highest values at the trailing edge. *Picea abies* showed the lowest predictions of occurrence and intensity of mortality in the core of the distribution and *Pinus pinaster* predicted mortality of both types was highest at the leading edge.

## 4 DISCUSSION

### 4.1 Drivers of occurrence and intensity of tree mortality across European tree ranges

Our results are in agreement with previous studies showing that the combination of drought and competition exacerbate the probability of mortality occurrence (Ruiz-Benito et al. 2013, Young, 2017) (Tables S11a, S12a), while intense events are driven by multiple factors (including drought and competition) (Anderegg et al., 2015; Jump et al. 2017; Seidl et al., 2017; Wood, Knapp, Muzika, Stambaugh, & Gu, 2018). As expected, the most intense mortality events were associated with low precipitation and high temperatures, the latter only for half of the species (Tables S11b, S12b). This may be explained by the synergic effects of low precipitation and warm temperatures with insect outbreaks (not included in our models) that may end into die-off events (Anderegg et al., 2015; Kurz et al., 2008; McDowell et al., 2011; Wood et al., 2018).

The importance of drought for the occurrence of mortality was higher in marginal populations that experienced drier than average conditions during the study period, suggesting that these populations are the most vulnerable to climate warming (Figure 1a-f and S6a-c). Although in exceptional cases, the highest level of relative drought was more detrimental for core than marginal populations (Figure S5a-d). Interestingly, high temperatures at the leading edge were correlated with the most intense events of mortality for many species, whereas the combination of drought and climatic marginality was less important in mortality intensity than occurrence (Figure 2d-f and S7b-c). Other studies have already observed an increase in background mortality in mild climates and at the northern margin of species ranges associated with warm temperature (Ruiz Benito et al. 2013; Neuman et al. 2017). The less important role of drought in explaining intensity of mortality could be explained by the time lag between stressful conditions and mortality responses, making it complicated to detect such small-time scale effects (Jump et al., 2017). We found a larger effect of competition on intensity of mortality at the trailing edge than at the core in temperate species (Figure 2a-c and S7a), which could be explained by the presence of more competitive and less prone to hydraulic risk Mediterranean species reaching their leading edge (Benito Garzón 2013, 2018).

### 4.2 Placing tree mortality at large geographical gradients

Mortality is an important component of demography and as such, its geographical patterns can be used to describe demographic and ecological differences of the core versus the peripheral populations (Purves, 2009; Pironon et al., 2017). We showed that the occurrence of mortality is greater in climatically marginal regions than at the climatic core of the species ranges. Furthermore, plots containing largest trees (higher DBH classes) and trees with the lowest growth rate (lower meanBAIj) were often associated with high mortality occurrence probability, suggesting that oldest trees are more likely to die (Vanoni et al., 2019; Hülsmann et al., 2017; Cailleret et al., 2017) (Table S11a).

The predicted probability of mortality occurrence was the highest in the trailing edge for most temperate species and the lowest in the leading edge for half of the Mediterranean species (Figure 3 and S4a). This suggest that the Mediterranean-temperate ecotone is a hotspot of forest composition changes, as previously suggested on the European gradients of water availability (Ruiz-Benito et al. 2017). Furthermore, in Mediterranean species, mortality occurrence was more likely to be higher in the trailing edge than in the core under intense drought (Figure 1d-f), suggesting that the southern part of the species ranges can be delimited by drought-induced mortality (Benito-Garzón et al., 2013; Benito Garzón et al., 2018; Gárate-Escamilla, Hampe, Vizcaíno-Palomar, Robson, & Benito Garzón, 2019; Kunstler et al., 2016). This result also highlights that drought increases in southern Europe are boosting background mortality in the last years (Benito Garzón et al., 2018; Carnicer et al., 2011; Druckenbrod et al., 2019; Neumann et al., 2017; Ruiz-Benito et al., 2013).

Conversely, the most intense events of mortality are evenly distributed in the European extent studied (Allen et al., 2015; Allen et al., 2010; Jump et al., 2009; Jump et al., 2017) and they are not related with old trees (DBH classes) or slow growth (meanBAI_j_) (Table S11b), suggesting they can affect all class of trees age and size as expected for die-off events. As such, die-off mortality can be affected by other important factors such as large competition (Young et al. 2017), pest emergence, fires etc. and result in unexpected demographic patterns. (Jump et al., 2017).

### 4.3 Limitations and perspectives

Although recently managed forests have been removed from our analysis according to management information in forest inventories (Ruiz-Benito et al. 2020), further legacy effects for which no information is available could affect mortality (Clark, Bell, Hersh, & Nichols, 2011; Csilléry et al., 2013, Young 2017). In addition, disturbance magnitude and duration could have contrasting lag effects and the differences in the time lag between surveys in our data can result in large uncertainties (Druckenbrod et al., 2019).

Our findings provide a new perspective to study tree mortality in forests from NFI data as both demographic and stochastic processes are exacerbated to some extent by drought stress but also by an interaction of climatic drivers that change across species ranges.

## Acknowledgements

This study was funded by the “Investments for the Future” program IdEx Bordeaux (ANR-10-IDEX-03-02) and the ATHENEE project (Nouvelle-Aquitaine Région). We thank the FunDivEUROPE project (European Union Seventh Framework Programme (FP7/2007-2013) under grant agreement no 265171) for the previous harmonization of the National Forest Inventory data used in this study. MAZ and PRB were supported by grant DARE; RTI2018-096884-B-C32 (MICINN, Spain).

