## Supplemental Tables for "Occurrence but not intensity of mortality rises towards the climatic trailing edge of tree species ranges in European forests"

### Supplementary Information

#### National Forest Inventories harmonization

The sample plot design was different among countries. For instance, the Spanish National Inventory recorded single sample plots in a 1 km by 1 km grid whereas the Finnish National Inventory followed a cluster design, with number and grid size depending on location while the German NFI used a 4 x 4 km quadrangle grid where the samples lied on the intersection points. Most NFIs followed a nested circular subplot design whose radius differs among NFI and within which trees of different size classes were monitored. The main differences among inventories are summarized in Table S1.

Table S1: Summary of information of the NFI design for each country: Belgium (Wallonia), Finland, France, Germany, Spain and Sweden Inventories. We included the sampling dates; plot type: permanent plots (PP) the years indicate the two campaigns used in the analysis, for temporary plots (TP) the years used in the analysis are indicated.  Grid size: indicates the grid dimension in km for each country. Distance between plots: indicates the distance between the plots within the grid. Plot radius: indicates the different radius (m) used within plots to sample trees. Sample tree DBH threshold: indicates the minimum DBH of the trees selected to sample a tree within a plot. N plot: number of plots per country. N trees: number of trees per country

| Country | Belgium-Wallonia | Finland | France | Germany | Spain | Sweden |
|---|---|---|---|---|---|---|
| Sampling dates and plot type | 1994-2003<br>2008-2011<br>PP | 1985-1986<br>1995<br>PP | 2005, 2006, 2007, 2008, 2009, 2010, 2011, 2012, 2013 and-2014<br>TP | 1986- 1990, West Germany only 2001-2002 (West and East Germany) | 1986-1996<br>1997-2007<br>PP | 2005-2007<br>2008-2010<br>TP and PP |
| Grid size (km) | 1x0.5 | 16x16 or 24x32 depending on the location. | 1x1 | 4x4, 2.83x2.83 or 2x2 depending on region | 1x1 | Vary |
| Sampling design (Distance between plots) | Single sample plots | Cluster design (100 or 300) | Single sample plots | Cluster design (150) | Single sample plots | Cluster design<br>(vary) |
| Radius (m) | 2.25, 4.5, 9, 12, 18 | 5.64, 9.77 | 6, 9, 15 | 1, 2, 5, 10, 25 | 5, 10, 15, 25 | 3.5, 10 |

| | | | | | | |
|---|---|---|---|---|---|---|
| Sample tree DBH threshold (cm) | 6.4 | 0 | 7.5 | **10** (1st)<br>7 (2nd) | 7.5 | 1 |
| N Plots | 1238 | 2487 | 60782 | 29914 | 48133 | 11338 |
| N Trees | 16011 | 39263 | 637830 | 295029 | 813464 | 187561 |

Table S2: List of species used for modeling tree mortality, number of plots in each species ranges defined (i.e. core, transition or marginal ranges, the latest divided in leading and trailing edges) and thresholds used to define the margins. Species: species name. Code: code used for each species. E= Ecology of the species (T=temperate; M=Mediterranean/Warm Temperate). *Castanea sativa* and *Pinus nigra* were included as Mediterranean species although they can also be considered as warm temperate species (San-Miguel-Ayanz, de Rigo, Caudullo, Houston Durrant, & Mauri, 2016).

NTrees: number of trees. NdeadTrees: number of dead trees. NPlots: number of plots. MT: margin thresholds used for the WPCA. CT: core thresholds used for the WPCA. NC: number of plots containing the species within the core of the species range. TE: Number of plots containing the species within the trailing edge of the species range. LE: number of plots containing the species within the leading edges of the species range. NT: Number of plots containing the species within the transition zone of the species range. NA: Number of plots containing the species but located outside of its known range. Min: Minimum census interval observed on all the plots in years. Mean: Average census interval observed on all the plots in years. Max: Maximum census interval observed on all the plots in years.

| Species | **E** | Code | NTrees | NdeadTrees (and %) | NPlots | MT | CT | NC | NTE | NLE | NT | NA | Min | Mean | Max |
|---|---|---|---|---|---|---|---|---|---|---|---|---|---|---|---|
| *Pinus sylvestris* L. | T | PIN SYL | 313266 | 16929 (5.40%) | 32223 | 0.8 | 0.6 | 11485 | 8480 | 582 | 7934 | 3742 | 4 | 9.7 | 20 |
| *Picea abies* (L.) H.Karst. | T | PIC ABI | 259151 | 7117 (2.74%) | 30639 | 0.8 | 0.6 | 6643 | 6264 | 1190 | 1495 | 15047 | 4 | 10.4 | 16 |
| *Pinus pinaster* Aiton. | M | PIN PINA | 177972 | 28834 (16.20) | 12610 | 0.8 | 0.6 | 5998 | 746 | 1357 | 2149 | 2360 | 7 | 11.3 | 16 |
| *Fagus sylvatica* L. | T | FAG SYL | 135856 | 3535 (2.60%) | 26237 | 0.7 | 0.6 | 20167 | 544 | 3872 | 1607 | 47 | 4 | 12.4 | 16 |
| *Pinus nigra* J.F.Arnold. | M | PIN NIG | 88427 | 4788 (5.41%) | 8360 | 0.7 | 0.6 | 4462 | 477 | 96 | 542 | 2783 | 4 | 11.2 | 20 |
| *Pinus halepensis* Mill. | M | PIN HAL | 87461 | 4929 (5.64%) | 9160 | 0.7 | 0.6 | 7110 | 606 | 411 | 867 | 166 | 2 | 11.3 | 19 |
| *Quercus robur* L. | T | QUE ROB | 85192 | 5483 (6,44%) | 21092 | 0.8 | 0.6 | 9750 | 3619 | 934 | 6778 | 11 | 9 | 12 | 16 |
| *Quercus ilex* L. | M | QUEILE | 84183 | 3543 (4.20 %) | 15267 | 0.8 | 0.6 | 9055 | 1839 | 646 | 3701 | 26 | 6 | 11.2 | 20 |

| | | | | | | | | | | | | | | | |
|---|---|---|---|---|---|---|---|---|---|---|---|---|---|---|---|
| *Quercus petraea* Liebl. | T | QUE PET | 77022 | 4071 (5.29%) | 17084 | 0.7 | 0.6 | 15700 | 486 | 250 | 617 | 31 | 9 | 12.8 | 20 |
| *Castanea sativa* Mill. | M | CAS SAT | 56235 | 12418 (22.08%) | 8683 | 0.7 | 0.6 | 6922 | 600 | 178 | 926 | 57 | 7 | 11 | 16 |
| *Abies alba* Mill. | T | ABI ALB | 50731 | 2517 (4.96%) | 8485 | 0.7 | 0.6 | 5635 | 511 | 1088 | 530 | 721 | 7 | 13.4 | 16 |
| *Quercus pubescens* Willd. | T | QUE PUB | 43767 | 2897 (6.62%) | 8406 | 0.7 | 0.5 | 7861 | 11 | 217 | 312 | 5 | 10 | 11.3 | 19 |
| *Quercus pyrenaica* Willd. | M | QUE PYR | 33157 | 1734 (5.23%) | 3835 | 0.7 | 0.6 | 2571 | 438 | 141 | 594 | 91 | 6 | 10.9 | 16 |
| *Fraxinus excelsior* L. | T | FRA EXC | 29899 | 888 (2.97%) | 9570 | 0.7 | 0.6 | 6537 | 1155 | 854 | 999 | 25 | 4 | 12.8 | 16 |
| *Pinus pinea* L. | M | PIN PIN | 27534 | 2438 (8.85%) | 3077 | 0.7 | 0.6 | 1516 | 246 | 282 | 667 | 366 | 6 | 10.9 | 14 |
| *Quercus suber* L. | M | QUE SUB | 25157 | 1143 (4.54%) | 3416 | 0.7 | 0.6 | 2396 | 461 | 172 | 351 | 36 | 6 | 11.4 | 16 |
| *Betula pendula* Roth. | T | BET PEN | 24222 | 1851(7.64%) | 8936 | 0.8 | 0.6 | 3843 | 1606 | 499 | 2517 | 471 | 4 | 11.6 | 16 |
| *Alnus glutinosa* (L). Gaertn. | T | ALB GLU | 14008 | 1067 (7.62%) | 2879 | 0.6 | 0.5 | 1658 | 780 | 109 | 316 | 16 | 4 | 9.8 | 16 |
| *Populus tremula* L. | T | POP TRE | 11739 | 1276 (10.87%) | 3903 | 0.7 | 0.6 | 2122 | 872 | 84 | 816 | 9 | 4 | 7.7 | 16 |
| *Acer pseudoplatanus* L. | T | ACE PSE | 10500 | 248 (2.36%) | 4442 | 0.55 | 0.5 | 2387 | 124 | 1284 | 220 | 427 | 5 | 13.5 | 16 |
| *Larix decidua* Mill. | T | LAR DEC | 10115 | 388 (3.84%) | 2768 | 0.6 | 0.5 | 315 | 1 | 159 | 73 | 2220 | 9 | 13.7 | 16 |
| *Populus nigra* L. | T | POP NIG | 4256 | 989 (23.24%) | 878 | 0.7 | 0.6 | 497 | 90 | 24 | 58 | 209 | 7 | 12.1 | 15 |

Table S3: Response variables analysed. Response variables are calculated for the $k^{th}$ individual of the $i^{th}$ individual plot of the $j^{th}$ species. Response: Response description and unit. Distribution:

distribution used to model the corresponding response. Calculation: final form of the response variable for modeling. Link: Link function used in the corresponding model.

| Response | Distribution | Calculation | Link |
|---|---|---|---|
| Species mortality occurrence (presence/absence at the plot level) | Binomial | $Y1_{ij}$ = 0 if no event recorded<br>$Y1_{ij}$ = 1 if at least one event recorded | logit |
| Species mortality intensity (Number of trees per ha per plot per year) | Zero-truncated negative binomial (NB) | $Y2_{ij} = \frac{dead\ trees_{ijk}}{numberof\ trees_{ijk}} \times \frac{1}{Years_i}$ | log |

Table S4: 21 climatic variables averaged over the last 30 years before the first inventory drought-related variables derived from SPEI indices calculated for the *i*th individual plot and climatic marginality categories obtained after the WPCA and DPCA (categories are calculated for the $i^{th}$ individual plot).We include the variable description/name and units/time period..

| Description | Unit/Time period |
|---|---|
| Annual mean temperature | °C |
| Mean diurnal temperature range | °C |
| Maximal temperature of the warmest month | °C |
| Minimal temperature of the coldest month | °C |
| Winter mean temperature | °C |
| Spring mean temperature | °C |
| Summer mean temperature | °C |
| Autumn mean temperature | °C |
| Annual precipitation | mm |
| Precipitation of the wettest month | mm |
| Precipitation of the driest month | mm |
| Winter precipitation | mm |
| Spring precipitation | mm |
| Summer precipitation | mm |
| Autumn precipitation | mm |
| Annual potential evapotranspiration | mm |
| Minimal monthly potential evapotranspiration | mm |
| Maximal monthly potential evapotranspiration | mm |
| Annual water balance | mm |
| Minimal monthly water balance | mm |

| | |
|---|---|
| Maximal monthly water balance | mm |
| Monthly Standardised Precipitation-Evapotranspiration Index of the last 12 months | Averaged on the time period elapsed between the two-sampling procedure |
| Monthly Standardised Precipitation-Evapotranspiration Index of the last 12 months | Minimum value on the time period elapsed between the two sampling procedure |
| Climatic trailing edge | No unit |
| Climatic leading edge | No unit |

Table S5: Cumulative variance explained by the first two axes of the weighted principal component analysis (WPCA) for each species. Species: Code of the species. Variance explained: Variance explained by the two first axes of the WPCA. Max Variable: climatic variable with the highest contribution to the PCA. Max%: percentage of the variance of the PCA explained by the Max Variable.

| Species | Variance explained | **Max Variable** | **Max %** |
|---|---|---|---|
| QUEILE | 78.287 | Annual water balance | 5.69 |
| PINPINA | 81.222 | Maximal temperature of the warmest month | 5.68 |
| PINSYL | 83.609 | Annual precipitation | 5.33 |
| PICABI | 82.541 | Maximal temperature of the warmest month | 5.46 |
| FAGSYL | 71.532 | Annual water balance | 6.16 |
| PINHAL | 78.394 | Annual water balance | 5.65 |
| QUEROB | 80.754 | Annual precipitation | 5.58 |
| PINNIG | 79.938 | Annual water balance | 5.63 |
| QUEPET | 75.681 | Annual water balance | 5.75 |
| CASSAT | 74.746 | Annual water balance | 5.9 |
| ABIALB | 81.14 | Annual water balance | 5.54 |
| QUEPYR | 79.25 | Maximal temperature of the warmest month | 5.78 |
| FRAEXC | 80.127 | Annual precipitation | 5.65 |
| PINPIN | 79.315 | Maximal temperature of the warmest month | 5.49 |
| QUESUB | 76.377 | Annual water balance | 5.84 |
| BETPEN | 82.326 | Annual precipitation | 5.64 |
| POPTRE | 83.335 | Annual water balance | 5.44 |
| POPNIG | 77.409 | Annual water balance | 6.03 |
| ACEPSE | 76.789 | Annual water balance | 5.79 |

| | | | |
|---|---|---|---|
| ALNGLU | 82.778 | Annual precipitation | 5.44 |
| LARDEC | 87.427 | Annual water balance | 5.22 |
| QUEPUB | 76.442 | Annual water balance | 5.91 |

Table S6: Biotic variables included in the model. Description: variable description. Calculation: equation used for variable calculation. Transformation: transformation of the variable before inclusion in the model. Abbreviation: abbreviated names of the variables as used in the manuscript.

| **Description** | **Calculation** | **Transformation** | **Abbreviation** |
|---|---|---|---|
| Number of years between surveys | $Years = Surveydate\,1_i - Surveydate\,2_i$ | None | yearsbetweenstudy |
| Species basal area increment ($ha^{-1}$ $yr^{-1}$) | $\sum \frac{Ba.ha2_{ijk} - Ba.ha1_{ijk}}{Years_i}$ | Square root | BAI |
| Mean species basal area increment of the plot ($ha^{-1}$ $yr^{-1}$) | $\frac{1}{N_{ijk}} \times \sum \frac{Ba.ha2_{ijk} - Ba.ha1_{ijk}}{Years_i}$ | Square root | meanBAIj |
| Mean diameter at breast height of the plot (cm) | $\frac{1}{N_{ij}} \times \sum dbh_{ijk}$ | Square root or logscaled | DBH (log or sqrt) |
| Number of trees in the plot (No. trees $ha^{-1}$ $yr^{-1}$) | $\sum k_{ij}$ | logscaled | Treenumber |
| Total basal area of the plot ($m^2$ $ha^{-1}$) | $\sum Ba.ha1_{ik}$ | Square root | BA |
| Total basal area of the same species ($m^2$ $ha^{-1}$) | $\sum Ba.ha1_{ijk}$ | Logscaled | BAcon |
| Total basal area of other species ($m^2$ $ha^{-1}$) | $\sum Ba.ha1_{ik} - \sum Ba.ha1_{ijk}$ | Square root | BAhetero |

Table S7: Biotic and climatic variables included in the best full model for each species. Species: code name of the species. Competition variable 1 and Competition variable2: competition related variables that were included in the model (see Table S3). Climatic variable 1 and Climatic variable 2: climate climatic related variables that were included in the full model among the 8 that were used (Table S5). DBH transformation indicates the type of the transformation applied to the DBH variable before it was included in the model: either log or square-rooted (sqrt) (Table S3)

| Species | Competition variable 1 | Competition variable 2 | Climatic variable 1 | Climatic variable 2 | DBH transformation |
|---|---|---|---|---|---|
| ABIALB | BA | BAhetero | Precipitation of the driest month | Winter mean temperature | log |
| ACEPSE | BA | BAcon | Precipitation of the driest month | Annual mean temperature | log |
| ALNGLU | BA | BAhetero | Precipitation of the driest | Annual mean | log |

| | | | | | |
|---|---|---|---|---|---|
| | | | month | temperature | |
| BETPEN | BA | BAcon | Precipitation of the driest month | Annual mean temperature | sqrt |
| CASSAT | BA | BAhetero | Precipitation of the driest month | Annual mean temperature | sqrt |
| FAGSYL | BA | BAcon | Precipitation of the direst month | Maximal temperature of the warmest month | sqrt |
| FRAEXC | BAcon | BAhetero | Annual precipitation | Maximal temperature of the warmest month | log |
| PICABI | BA | BAcon | Precipitation of the wettest month | Maximal temperature of the warmest month | sqrt |
| PINPINA | BA | BAhetero | Precipitation of the driest month | Annual mean temperature | log |
| PINHAL | BA | BAcon | Annual water balance | Maximal temperature of the warmest month | log |
| PINNIG | BA | BAcon | Annual precipitation | Maximal temperature of the warmest month | log |
| PINPIN | BA | BAcon | Precipitation of the driest month | Maximal temperature of the warmest month | log |
| PINSYL | BA | BAcon | Precipitation of the wettest month | Annual mean temperature | log |
| POPNIG | BA | BAcon | Annual precipitation | Maximal temperature of the warmest month | log |
| POPTRE | BA | BAhetero | Precipitation of the driest month | Annual mean temperature | log |
| QUEILE | BAcon | BAhetero | Annual precipitation | Annual mean temperature | sqrt |
| QUEPET | BA | BAcon | Annual precipitation | Annual mean temperature | sqrt |
| QUEPYR | BA | BAcon | Precipitation of the driest month | Annual mean temperature | sqrt |
| QUEROB | BA | BAhetero | Precipitation of the wettest month | Maximal temperature of the warmest month | sqrt |
| QUESUB | BA | BAcon | Annual precipitation | Annual mean temperature | sqrt |

As it is not possible to calculate a Variation Inflation Factor value for models including qualitative variables, we calculated the VIF for each quantitative variable included in the best predictive model for each species. In addition, to quantify the collinearity between climatic variables and the climatic marginality, we calculated the VIF for the marginality index, which is the climatic marginality score we obtained after weighting the PCA scores. In the next following tables, all the scores for each variable is reported.

Table S8a VIF values calculated on the variables included in the best predictive binomial model for each species. For each species, there is one column with the variable names that are included in the model and one column with the VIF scores calculated. Marginality index VIF's values are in bold.

| ABIALB Variables | VIF | ACEPSE Variables | VIF | ALNGLU Variables | VIF | BETPEN Variables | VIF | CASSAT Variables | VIF | FAGSYL Variables | VIF | FRAEXC Variables | VIF | PICABI Variables | VIF | PINHAL Variables | VIF | PINNIG Variables | VIF |
|---|---|---|---|---|---|---|---|---|---|---|---|---|---|---|---|---|---|---|---|
| BAIj | 9.42 | meanBAIj | 2.23 | BAIj | 7.1 | BAIj | 7.18 | BAIj | 6.36 | BAIj | 6.57 | BAIj | 9.02 | BAIj | 6.83 | meanBAIj | 1.89 | meanBAIj | 1.62 |
| meanBAIj | 9.47 | DBH | 3.73 | meanBAIj | 6.88 | meanBAIj | 7.85 | meanBAIj | 6.24 | meanBAIj | 6.91 | meanBAIj | 8.13 | meanBAIj | 5.5 | DBH | 3.43 | bio5 | 2.16 |
| DBH | 2.79 | treeNbr | 2.91 | DBH | 2.56 | DBH | 5.01 | DBH | 2.78 | treeNbr | 2.44 | DBH | 6.65 | DBH | 4.55 | treeNbr | 5.61 | bio12 | 2.47 |
| treeNbr | 2.72 | bio14 | 2.74 | treeNbr | 2.43 | treeNbr | 2.39 | treeNbr | 2.71 | min_spei12 | 1.67 | treeNbr | 3.01 | bio5 | 1.29 | yearsbetweensurveys | 3.63 | min_spei12 | 2.28 |
| BA | 3.93 | mean_spei12 | 1.56 | bio14 | 1.39 | bio1 | 1.84 | yearsbetweensurveys | 2.13 | mean_spei12 | 1.43 | Yearsbetweensurveys | 1.91 | bio13 | 4.36 | bio5 | 5.2 | mean_spei12 | 2.25 |
| BAhetero | 4.2 | BA | 3.02 | **Marginality** | **2.43** | min_spei12 | 1.58 | bio1 | 1.94 | BA | 1.85 | BAhetero | 2.33 | min_spei12 | 2.85 | ppet.mean | 7.44 | BA | 2.68 |
| **Marginality** | **1.18** | bio1 | 2.19 | min_spei12 | 1.7 | mean_spei12 | 1.45 | bio14 | 1.95 | BAcon | 2.31 | **Marginality** | **1.17** | BAcon | 10.9 | min_spei12 | 1.98 | BAcon | 2.95 |
| bio14 | 1.52 | min_spei12 | 1.59 | bio1 | 2.65 | BA | 2.47 | mean_spei12 | 1.83 | **Marginality** | **2.11** | mean_spei12 | 2.42 | **Marginality** | **4.37** | mean_spei12 | 1.67 | **Marginality** | **1.42** |
| min_spei12 | 2.07 | BAcon | 4.27 | BA | 4.58 | BAcon | 6.53 | BA | 4.47 | bio14 | 2.02 | bio5 | 1.3 | mean_spei12 | 2.53 | BA | 6.84 | - | - |
| tmean.djf | 1.32 | **Marginality** | **3.34** | BAhetero | 4.61 | **Marginality** | **1.58** | **Marginality** | **2.41** | bio5 | 1.47 | BAcon | 8.5 | BA | 3.76 | BAcon | 7.09 | - | - |
| mean_spei12 | 2.16 | - | - | mean_spei12 | 1.93 | - | - | BAhetero | 4.45 | - | - | min_spei12 | 2.34 | - | - | **Marginality** | **1.13** | - | - |
| - | - | - | - | - | - | - | - | min_spei12 | 2.12 | - | - | - | - | - | - | - | - | - | - |

| PINPINA Variables | VIF | PINPIN Variables | VIF | PINSYL Variables | VIF | POPNIG Variables | VIF | POPTRE Variables | VIF | QUEILE Variables | VIF | QUEPYR Variables | VIF | QUEPET Variables | VIF | QUEROB Variables | VIF | QUESUB Variables | VIF |
|---|---|---|---|---|---|---|---|---|---|---|---|---|---|---|---|---|---|---|---|

| | | | | | | | | | | | | | | | | | | | |
|---|---|---|---|---|---|---|---|---|---|---|---|---|---|---|---|---|---|---|---|
| meanBAIj | 2.36 | meanBAIj | 2.24 | meanBAIj | 1.82 | meanBAIj | 1.96 | BAIj | 5.72 | BAIj | 2.22 | meanBAIj | 1.8 | BAIj | 9.91 | BAIj | 5.39 | meanBAIj | 1.86 |
| DBH | 2.9 | DBH | 4.46 | DBH | 4.71 | DBH | 5.3 | meanBAIj | 6.83 | DBH | 2.8 | bio1 | 1.33 | meanBAIj | 8.16 | meanBAIj | 5.91 | DBH | 6.23 |
| treeNbr | 3.87 | bio5 | 3.96 | treeNbr | 2.09 | treeNbr | 2.15 | DBH | 1.96 | yearsbetweensurveys | 1.65 | bio14 | 2.85 | DBH | 4.89 | DBH | 2.9 | treeNbr | 4.93 |
| min_spei12 | 1.85 | bio14 | 5.47 | bio1 | 2.27 | bio5 | 1.68 | treeNbr | 2.56 | bio1 | 1.25 | mean_spei12 | 2.81 | treeNbr | 2.52 | treeNbr | 2.97 | yearsbetweensurveys | 2.37 |
| mean_spei12 | 2.45 | min_spei12 | 2.85 | bio13 | 1.15 | bio12 | 1.28 | bio14 | 1.55 | bio12 | 1.67 | BAcon | 2.07 | bio12 | 1.26 | yearsbetweensurveys | 2.36 | bio1 | 1.43 |
| **Marginality** | **1.4** | BA | 3.35 | BAcon | 8.7 | min_spei12 | 1.2 | mean_spei12 | 2.16 | min_spei12 | 1.79 | min_spei12 | 2.37 | min_spei12 | 1.43 | bio5 | 1.76 | bio12 | 1.21 |
| bio14 | 2.08 | BAcon | 6.18 | **Marginality** | **2.6** | BA | 3.03 | BA | 1.83 | BAcon | 2.93 | BA | 2.41 | mean_spei12 | 1.43 | bio13 | 2.77 | BAcon | 6.93 |
| bio1 | 1.32 | mean_spei12 | 1.93 | mean_spei12 | 2.27 | BAcon | 7.39 | bio1 | 2.13 | mean_spei12 | 1.81 | **Marginality** | **1.54** | BA | 2.24 | mean_spei12 | 2.9 | min_spei12 | 1.4 |
| BA | 4.51 | **Marginality** | **1.46** | min_spei12 | 2.69 | **Marginality** | **1.57** | min_spei12 | 2.15 | BAhetero | 1.84 | - | - | BAcon | 7.96 | BA | 5.47 | BA | 4.08 |
| BAhetero | 2.47 | - | - | BA | 3.68 | - | - | **Marginality** | **1.94** | **Marginality** | **1.25** | - | - | bio1 | 1.39 | BAhetero | 6.66 | mean_spei12 | 2.09 |
| - | - | - | - | - | - | - | - | - | - | - | - | - | - | - | - | min_spei12 | 2.7 | **Marginality** | **1.21** |
| - | - | - | - | - | - | - | - | - | - | - | - | - | - | - | - | **Marginality** | **3.08** | - | - |

Table S8b VIF values calculated on the variables included in the best predictive negative binomial model for each species. For each species, there is one column with the variable names that are included in the model and one column with the VIF scores calculated. Marginality index VIF's values are in bold.

| ABIALB Variables | VIF | ACEPSE Variables | VIF | ALNGLU Variables | VIF | BETPEN Variables | VIF | CASSAT Variables | VIF | FRAEXC Variables | VIF | FAGSYL Variables | VIF | PICABI Variables | VIF | PINHAL Variables | VIF | PINNIG Variables | VIF |
|---|---|---|---|---|---|---|---|---|---|---|---|---|---|---|---|---|---|---|---|
| BAIj | 9.17 | meanBAIj | 2.23 | BAIj | 7.1 | BAIj | 7.18 | BAIj | 5.79 | BAIj | 8.94 | BAIj | 6.44 | BAIj | 6.99 | meanBAIj | 1.92 | meanBAIj | 1.62 |
| meanBAIj | 9.36 | DBH | 3.73 | meanBAIj | 6.88 | meanBAIj | 7.85 | meanBAIj | 5.71 | meanBAIj | 8.06 | meanBAIj | 6.42 | meanBAIj | 5.77 | DBH | 3.61 | bio5 | 2.16 |
| DBH | 2.82 | treeNbr | 2.91 | DBH | 2.56 | DBH | 5.01 | DBH | 2.85 | DBH | 6.14 | treeNbr | 2.41 | DBH | 4.57 | treeNbr | 5.6 | bio12 | 2.47 |
| treeNbr | 2.69 | bio14 | 2.74 | treeNbr | 2.43 | treeNbr | 2.39 | treeNbr | 2.71 | treeNbr | 2.8 | min_spei12 | 1.69 | bio5 | 1.31 | yearsbetweensurveys | 3.63 | min_spei12 | 2.28 |
| BA | 3.92 | mean_spei12 | 1.56 | bio14 | 1.39 | bio1 | 1.84 | yearsbetweensurveys | 2.1 | yearsbetweensurveys | 1.87 | mean_spei12 | 1.46 | bio13 | 4.43 | bio5 | 5.02 | mean_spei12 | 2.25 |
| BAhetero | 4.18 | BA | 3.02 | **Marginality** | **2.43** | min_spei12 | 1.58 | bio1 | 1.96 | BAhetero | 2.32 | BA | 1.81 | min_spei12 | 2.93 | ppet.mean | 7.21 | BA | 2.68 |
| **Marginality** | **1.18** | bio1 | 2.19 | min_spei12 | 1.7 | mean_spei12 | 1.45 | bio14 | 1.98 | **Marginality** | **1.2** | BAcon | 2.35 | BAcon | 10.84 | min_spei12 | 1.93 | BAcon | 2.95 |
| bio14 | 1.52 | min_spei12 | 1.59 | bio1 | 2.65 | BA | 2.47 | mean_spei12 | 1.87 | mean_spei12 | 2.44 | **Marginality** | **2.24** | **Marginality** | **4.42** | mean_spei12 | 1.66 | **Marginality** | **1.42** |
| min_spei12 | 2.08 | BAcon | 4.27 | BA | 4.58 | BAcon | 6.53 | BA | 4.45 | bio5 | 1.31 | bio14 | 2.04 | mean_spei12 | 2.6 | BA | 6.81 | - | - |
| tmean.djf | 1.32 | **Marginality** | **3.34** | BAhetero | 4.61 | **Marginality** | **1.58** | **Marginality** | **2.42** | BAcon | 7.98 | bio5 | 1.54 | BA | 3.78 | BAcon | 7.27 | - | - |
| mean_spei12 | 2.16 | - | - | mean_spei12 | 1.93 | - | - | BAhetero | 4.39 | min_spei12 | 2.32 | - | - | - | - | **Marginality** | **1.13** | - | - |
| - | - | - | - | - | - | - | - | min_spei12 | 2.21 | - | - | - | - | - | - | - | - | - | - |

| PINPINA Variables | VIF | PINPIN Variables | VIF | PINSYL Variables | VIF | POPNIG Variables | VIF | POPTRE Variables | VIF | QUEILE Variables | VIF | QUEPET Variables | VIF | QUEROB Variables | VIF | QUEPYR Variables | VIF | QUESUB Variables | VIF |
|---|---|---|---|---|---|---|---|---|---|---|---|---|---|---|---|---|---|---|---|
| meanBAIj | 2.38 | meanBAIj | 2.24 | meanBAIj | 1.77 | meanBAIj | 1.96 | BAIj | 5.72 | BAIj | 2.18 | BAIj | 10.34 | BAIj | 5.34 | meanBAIj | 1.8 | meanBAIj | 1.86 |
| DBH | 2.91 | DBH | 4.46 | DBH | 4.39 | DBH | 5.3 | meanBAIj | 6.83 | DBH | 2.64 | meanBAIj | 8.56 | meanBAIj | 5.83 | bio1 | 1.33 | DBH | 6.23 |
| treeNbr | 3.83 | bio5 | 3.96 | treeNbr | 2.1 | treeNbr | 2.15 | DBH | 1.96 | yearsbetweensurveys | 1.61 | DBH | 5.04 | DBH | 2.84 | bio14 | 2.85 | treeNbr | 4.93 |

| | | | | | | | | | | | | | | | | | | | |
|---|---|---|---|---|---|---|---|---|---|---|---|---|---|---|---|---|---|---|---|
| min_spei12 | 1.92 | bio14 | 5.47 | bio1 | 2.39 | bio5 | 1.68 | treeNbr | 2.56 | bio1 | 1.25 | treeNbr | 2.6 | treeNbr | 2.94 | mean_spei12 | 2.81 | yearsbetweensurveys | 2.37 |
| mean_spei12 | 2.52 | min_spei12 | 2.85 | bio13 | 1.15 | bio12 | 1.28 | bio14 | 1.55 | bio12 | 1.66 | bio12 | 1.28 | yearsbetweensurveys | 2.42 | BAcon | 2.07 | bio1 | 1.43 |
| **Marginality** | **1.39** | BA | 3.35 | BAcon | 8.7 | min_spei12 | 1.2 | mean_spei12 | 2.16 | min_spei12 | 1.8 | min_spei12 | 1.44 | bio5 | 1.78 | min_spei12 | 2.37 | bio12 | 1.21 |
| bio14 | 2.22 | BAcon | 6.18 | **Marginality** | **2.72** | BA | 3.03 | BA | 1.83 | BAcon | 2.89 | mean_spei12 | 1.43 | bio13 | 2.8 | BA | 2.41 | BAcon | 6.93 |
| bio1 | 1.32 | mean_spei12 | 1.93 | mean_spei12 | 2.29 | BAcon | 7.39 | bio1 | 2.13 | mean_spei12 | 1.73 | BA | 2.29 | mean_spei12 | 2.86 | **Marginality** | **1.54** | min_spei12 | 1.4 |
| BA | 4.41 | **Marginality** | **1.46** | min_spei12 | 2.64 | **Marginality** | **1.57** | min_spei12 | 2.15 | BAhetero | 1.77 | BAcon | 8.22 | BA | 5.58 | - | - | BA | 4.08 |
| BAhetero | 2.4 | - | - | BA | 3.64 | - | - | **Marginality** | **1.94** | **Marginality** | **1.23** | bio1 | 1.45 | BAhetero | 6.85 | - | - | mean_spei12 | 2.09 |
| - | - | - | - | - | - | - | - | - | - | - | - | - | - | min_spei12 | 2.69 | - | - | **Marginality** | **1.21** |
| - | - | - | - | - | - | - | - | - | - | - | - | - | - | **Marginality** | **3.11** | - | - | - | - |

Table S9: Predicted mortality quartiles by species. Species: Code of the species. Q1BIN: value of the first predicted occurrence of mortality quartile. Q3BIN: value of the third predicted occurrence of mortality quartile. Q1ZTNB: value of the first predicted intensity of mortality quartile (in ‰ of tree per hectare per year). Q3ZTNB: value of the third predicted intensity of mortality quartile (in ‰ of tree per hectare per year). NA: no model was fitted.

| Species | Q1BIN | Q3BIN | Q1ZTNB | Q3ZTNB |
|---|---|---|---|---|
| PINPINA | **0.12** | **0.57** | 22 | 39 |
| ABIALB | 0.033 | 0.2 | 22 | 44 |
| ACEPSE | 0.0049 | 0.03 | 21 | 61 |
| ALNGLU | 0.045 | 0.23 | 26 | 56 |
| BETPEN | 0.017 | 0.11 | 27 | 64 |
| CASSAT | **0.12** | **0.47** | 34 | 64 |
| FAGSYL | 0.014 | 0.072 | 18 | 31 |
| FRAEXC | 0.011 | 0.055 | 23 | 54 |
| PICABI | 0.036 | 0.19 | 18 | 35 |
| PINHAL | 0.035 | 0.22 | 19 | 33 |
| PINNIG | 0.028 | 0.17 | 18 | 31 |
| PINPIN | 0.038 | 0.24 | 23 | 40 |
| PINSYL | 0.048 | 0.25 | 18 | 35 |
| POPNIG | 0.026 | 0.24 | NA | NA |
| POPTRE | 0.027 | 0.15 | **40** | **70** |
| QUEILE | 0.026 | 0.11 | 20 | 37 |
| QUEPET | 0.025 | 0.13 | 27 | 48 |
| QUEPYR | 0.036 | 0.2 | 21 | 38 |
| QUEROB | 0.027 | 0.15 | 28 | 56 |
| QUESUB | 0.052 | 0.2 | 17 | 33 |

Table S10: Observed occurrence and intensity of mortality by plot. Code: Code used for each species. Obs plot mortality: Average intensity of mortality = percentage of dead trees by year and by hectare based on all plot. Obs plot mortality (occurrence): Average intensity of mortality in plots where mortality occurs only= percentage of dead trees by year and by hectare. Minimum and maximum values are in parenthesis for each column.

| Species | Obs plot mortality<br>(mean% dead trees/yr/ha) | Obs plot mortality<br>(occurrence)<br>(mean% dead trees/yr/ha) |
|---|---|---|
| ABIALB | 0.61<br>(0 - 20) | 4.4<br>(0.1 - 20) |

| | | |
|---|---|---|
| ACEPSE | **0.21**<br>(0 - 20) | 6.46<br>(0.3 - 20) |
| ALNGLU | 1.15<br>(0 - 25) | 6.66<br>(0.2 - 25) |
| BETPEN | 1.15<br>(0 - 25) | 9.68<br>(0.2 - 25) |
| CASSAT | **2.48**<br>(0 - 20) | 7.17<br>(0.1 - 20) |
| FAGSYL | 0.23<br>(0 - 20) | 3.65<br>(0.1 - 20) |
| FRAEXC | 0.36<br>(0 - 20) | 6.53<br>(0.1 - 20) |
| PICABI | 0.47<br>(0 - 25) | 3.81<br>(0.1 - 25) |
| PINPINA | 1.71<br>(0 - 20) | 4.4<br>(0.1 - 20) |
| PINHAL | 0.51<br>(0 - 20) | 3.52<br>(0.1 - 20) |
| PINNIG | 0.36<br>(0 - 10.6) | **2.95**<br>(0.1 - 10.6) |
| PINPIN | 0.79<br>(0 - 10) | 4.43<br>(0.1 - 10) |
| PINSYL | 0.67<br>(0 - 25) | 3.94<br>(0.1 - 25) |
| POPNIG | 1.5<br>(0 - 20) | 7.35<br>(0.6 - 20) |
| POPTRE | 1.67<br>(0 - 25) | **10.73**<br>(0.2 - 25) |
| QUEILE | 0.36<br>(0 - 20) | 4.23<br>(0.1 - 20) |
| QUEPET | 0.58<br>(0 - 20) | 5.08<br>(0.1 - 20) |
| QUEPYR | 0.56<br>(0 - 20) | 4.07<br>(0.1 - 20) |
| QUEROB | 0.67<br>(0 - 20) | 5.49<br>(0.1 - 20) |
| QUESUB | 0.51<br>(0 - 20) | 3.52<br>(0.1 - 20) |

Table S11a The five simple effects with the most important coefficient estimated by GLMM for each species. Species: Code of the species name. EV: Variable acronym with the importance number indicated. EV t-value: t-value associated with the variable. SE: Standard error associated with the variable.

| Species | EV1<br>EV1 t-value<br>EV1 SE | EV2<br>EV2 t-value<br>EV2 SE | EV3<br>EV3 t-value<br>EV3 SE | EV4<br>EV4 t-value<br>EV4 SE | EV5<br>EV5 t-value<br>EV5 SE |
|---|---|---|---|---|---|
| CASSAT | DBH<br>21.194<br>0.039 | treeNbr<br>13.694<br>0.007 | mean_spei12<br>-10.989<br>0.109 | meanBAIj<br>-10.528<br>0.134 | BAIj<br>10.229<br>0.101 |
| PINHAL | meanBAIj<br>-15.674<br>0.072 | BAcon<br>11.139<br>0.133 | ppet.mean<br>8.116<br>0.284 | (Intercept)<br>-7.429<br>0.542 | min_spei12<br>-5.171<br>0.641 |
| PINNIG | meanBAIj<br>-11.254<br>0.101 | BAcon<br>8.926<br>0.349 | BA<br>-5.075<br>0.327 | bio5<br>3.637<br>0.449 | I(min_spei12^2)<br>3.433<br>0.364 |
| PINPINA | DBH<br>23.161<br>0.048 | meanBAIj<br>-15.51<br>0.085 | treeNbr<br>13.381<br>0.003 | (Intercept)<br>-8.581<br>0.868 | min_spei12<br>-6.6<br>1.064 |
| PINPIN | meanBAIj<br>-8.747<br>0.136 | BAcon<br>7.43<br>0.257 | bio14<br>4.219<br>0.345 | I(min_spei12^2)<br>3.394<br>0.383 | I(bio5^2)<br>3.252<br>0.303 |
| QUEILE | BAcon<br>19.442<br>0.087 | BAIj<br>-10.681<br>0.057 | bio1<br>7.851<br>0.061 | bio12<br>5.482<br>0.222 | DBH<br>-5.4<br>0.069 |
| QUEPYR | (Intercept)<br>-22.406<br>0.114 | BAcon<br>12.968<br>0.139 | meanBAIj<br>-8.08<br>0.112 | bio1<br>4.351<br>0.091 | bio14<br>-4.289<br>0.449 |
| QUESUB | meanBAIj<br>-8.253 | (Intercept)<br>-5.306 | DBH<br>4.59 | BAcon<br>3.884 | treeNbr<br>3.45 |

| | | | | | |
|---|---|---|---|---|---|
| | 0.088 | 0.4 | 0.17 | 0.191 | 0.012 |
| ABIALB | DBH<br>13.627<br>0.091 | meanBAIj<br>-9.734<br>0.22 | BAIj<br>7.466<br>0.182 | BA<br>-6.447<br>0.098 | treeNbr<br>5.103<br>0.014 |
| ACEPSE | treeNbr<br>7.165<br>0.029 | DBH<br>6.869<br>0.183 | (Intercept)<br>-5.751<br>1.41 | meanBAIj<br>-2.886<br>0.171 | mean_spei12<br>-2.565<br>0.415 |
| ALNGLU | DBH<br>9.591<br>0.104 | (Intercept)<br>-7.349<br>0.502 | meanBAIj<br>-6.271<br>0.291 | BAIj<br>4.467<br>0.223 | treeNbr<br>2.577<br>0.016 |
| BETPEN | DBH<br>8.319<br>0.128 | treeNbr<br>7.349<br>0.017 | meanBAIj<br>-6.3<br>0.322 | bio1<br>4.839<br>0.226 | mean_spei12<br>-4.369<br>0.371 |
| FAGSYL | BAcon<br>26.37<br>0.065 | (Intercept)<br>-14.291<br>0.277 | BA<br>-12.058<br>0.061 | mean_spei12<br>-10.574<br>0.139 | meanBAIj<br>-8.507<br>0.129 |
| FRAEXC | DBH<br>12.397<br>0.088 | treeNbr<br>8.091<br>0.016 | BAhetero<br>-6.265<br>0.081 | meanBAIj<br>-5.796<br>0.259 | (Intercept)<br>-4.44<br>2.035 |
| PICABI | BAcon<br>12.623<br>0.139 | meanBAIj<br>-10.222<br>0.098 | (Intercept)<br>-8.77<br>0.359 | BAIj<br>4.238<br>0.091 | Plotcat2<br>4.22<br>0.37 |
| PINSYL | meanBAIj<br>-19.998<br>0.037 | treeNbr<br>11.48<br>0.003 | (Intercept)<br>-10.25<br>0.274 | bio13<br>8.881<br>0.115 | bio1<br>8.019<br>0.055 |
| POPNIG | treeNbr<br>5.066<br>0.041 | (Intercept)<br>-4.098<br>1.407 | DBH<br>3.466<br>0.342 | meanBAIj<br>-3.302<br>0.252 | NA<br>NA<br>NA |
| POPTRE | DBH<br>11.125 | mean_spei12<br>-7.464 | (Intercept)<br>-5.993 | meanBAIj<br>-5.692 | treeNbr<br>4.825 |

| | | | | | |
|---|---|---|---|---|---|
| | 0.103 | 0.298 | 0.737 | 0.299 | 0.014 |
| QUEROB | meanBAIj<br>-15.942<br>0.147 | DBH<br>14.73<br>0.063 | BAIj<br>13.196<br>0.101 | treeNbr<br>8.433<br>0.009 | mean_spei12<br>-8.427<br>0.181 |

Table S11b. The five simple effects with the most important coefficient estimated by the ZTNBMM for each species. Species: Code for the species name. EV: Variable acronym with the importance number indicated. EV t-value: t-value associated with the variable. SE: Standard error associated with the variable.

| Species | EV1<br>EV1 t-value<br>EV 1 SE | EV2<br>EV2 t-value<br>EV 2 SE | EV3<br>EV3 t-value<br>EV 3 SE | EV4<br>EV4 t-value<br>EV 4 SE | EV5<br>EV5 t-value<br>EV 5 SE |
|---|---|---|---|---|---|
| CASSAT | meanBAIj<br>-18.706<br>0.024 | BA<br>-11.518<br>0.029 | DBH<br>9.201<br>0.026 | BAhetero<br>6.536<br>0.055 | treeNbr<br>6.06<br>0.003 |
| PINHAL | (Intercept)<br>25.362<br>0.169 | BAIj<br>-17.034<br>0.031 | yearsbetweensurveys<br>-8.12<br>0.017 | treeNbr<br>7.473<br>0.003 | DBH<br>4.731<br>0.035 |
| PINNIG | (Intercept)<br>8.825<br>0.603 | BAIj<br>-5.784<br>0.067 | mean_spei12<br>-3.947<br>0.965 | yearsbetweensurveys<br>-3.35<br>0.044 | I(min_spei12^2)<br>2.562<br>0.276 |
| PINPINA | (Intercept)<br>25.819<br>0.182 | BAIj<br>-10.569<br>0.049 | yearsbetweensurveys<br>-10.202<br>0.015 | treeNbr<br>8.749<br>0.003 | DBH<br>4.008<br>0.035 |
| PINPIN | BAIj<br>-9.19<br>0.062 | (Intercept)<br>7.776<br>0.597 | BAcon<br>5.653<br>0.174 | DBH<br>-4.655<br>0.119 | BA<br>-3.773<br>0.092 |
| QUEILE | BAIj<br>-12.383 | (Intercept)<br>9.722 | treeNbr<br>6.538 | yearsbetweensurveys<br>-4.872 | mean_spei12<br>-3.051 |

| | | | | | |
|---|---|---|---|---|---|
| | 0.032 | 0.447 | 0.004 | 0.031 | 0.492 |
| QUEPYR | (Intercept)<br>16.028<br>0.333 | BAIj<br>-8.706<br>0.054 | yearsbetweensurveys<br>-5.803<br>0.032 | I(mean_spei12^2)<br>-2.943<br>0.936 | mean_spei12<br>2.806<br>1.357 |
| QUESUB | (Intercept)<br>19.853<br>0.227 | yearsbetweensurveys<br>-7.127<br>0.02 | BAIj<br>-6.327<br>0.044 | I(bio1^2)<br>3.859<br>0.029 | BAcon<br>-3.839<br>0.064 |
| ABIALB | BAIj<br>-11.515<br>0.03 | (Intercept)<br>5.885<br>0.665 | BAhetero<br>3.922<br>0.048 | mean_spei12<br>2.861<br>0.215 | I(mean_spei12^2)<br>-2.814<br>0.227 |
| ACEPSE | Plotcat2<br>-4.927<br>0.393 | BAIj<br>-4.531<br>0.176 | BAcon<br>-4.503<br>0.146 | (Intercept)<br>4.337<br>1.018 | meanBAIj<br>3.4<br>0.237 |
| ALNGLU | (Intercept)<br>19.878<br>0.198 | meanBAIj<br>-7.362<br>0.06 | BAhetero<br>3.263<br>0.095 | BA<br>-3.207<br>0.067 | treeNbr<br>-2.736<br>0.006 |
| BETPEN | (Intercept)<br>13.971<br>0.33 | BAcon<br>-10.473<br>0.065 | DBH<br>7.938<br>0.053 | meanBAIj<br>-7.384<br>0.058 | I(min_spei12^2)<br>4.331<br>0.111 |
| FAGSYL | (Intercept)<br>10.725<br>0.369 | BAcon<br>-7.508<br>0.064 | BAIj<br>-5.695<br>0.027 | DBH<br>4.018<br>0.039 | mean_spei12<br>3.925<br>0.095 |
| FRAEXC | (Intercept)<br>32.717<br>0.126 | BAcon<br>-13.558<br>0.053 | DBH<br>8.436<br>0.064 | bio12<br>-4.957<br>0.094 | BAhetero<br>4.361<br>0.097 |
| PICABI | BAIj<br>-16.042<br>0.021 | (Intercept)<br>11.545<br>0.313 | treeNbr<br>-6.722<br>0.002 | DBH<br>-5.825<br>0.026 | Plotcat1<br>4.998<br>0.119 |
| PINSYL | BAIj<br>-14.861 | (Intercept)<br>13.979 | BAcon<br>-10.939 | treeNbr<br>9.661 | yearsbetweensurveys<br>-5.742 |

| | | | | | |
|---|---|---|---|---|---|
| | 0.019 | 0.297 | 0.064 | 0.002 | 0.019 |
| POPTRE | (Intercept)<br>32.606<br>0.129 | BAIj<br>-8.301<br>0.046 | DBH<br>4.879<br>0.056 | BAhetero<br>4.453<br>0.059 | yearsbetweensurveys<br>-2.663<br>0.021 |
| QUEPET | BAcon<br>-17.943<br>0.049 | DBH<br>14.768<br>0.039 | (Intercept)<br>6.703<br>0.775 | min_spei12<br>6.294<br>0.142 | bio12<br>-5.39<br>0.037 |
| QUEROB | (Intercept)<br>11.161<br>0.406 | BAIj<br>-9.301<br>0.048 | min_spei12<br>4.418<br>0.152 | I(bio5^2)<br>3.259<br>0.029 | I(min_spei12^2)<br>3.229<br>0.07 |

Table S11c: The five interaction effects with the most important coefficient estimated by the GLMM for each species. Species: Code of the species name. EV: Variable acronym with the importance number indicated. EV t-value: t-value associated with the variable. SE: Standard error associated with the variable.

| Species | EV1<br>EV1 t-value<br>EV 1 SE | EV2<br>EV2 t-value<br>EV 2 SE | EV3<br>EV3 t-value<br>EV 3 SE | EV4<br>EV4 t-value<br>EV 4 SE | EV5<br>EV5 t-value<br>EV 5 SE |
|---|---|---|---|---|---|
| CASSAT | bio14:min_spei12<br>4.672<br>0.139 | bio14:mean_spei12<br>-4.51<br>0.188 | bio14:BA<br>3.523<br>0.033 | min_spei12:Plotcat1<br>3.401<br>0.5 | mean_spei12:Plotcat1<br>-2.724<br>0.483 |
| PINHAL | ppet.mean:min_spei12<br>6.745<br>0.194 | bio5:mean_spei12<br>6.239<br>0.225 | BA:BAcon<br>-4.957<br>0.042 | min_spei12:mean_spei12<br>3.291<br>0.445 | min_spei12:Plotcat2<br>-3.173<br>0.704 |
| PINNIG | min_spei12:BAcon<br>3.725<br>0.231 | min_spei12:BA<br>-3.596<br>0.22 | BA:Plotcat1<br>3.559<br>0.263 | bio5:BA<br>-3.479<br>0.082 | mean_spei12:Plotcat2<br>-3.196<br>1.668 |
| PINPINA | min_spei12:bio1<br>-6.537 | min_spei12:Plotcat1<br>4.968 | Plotcat2:BAhetero<br>-3.726 | mean_spei12:bio1<br>3.538 | mean_spei12:BAhetero<br>3.044 |

| | | | | | |
|---|---|---|---|---|---|
| | 0.036 | 0.756 | 0.106 | 0.185 | 0.126 |
| PINPIN | bio5:bio14<br>3.3<br>0.552 | bio5:Plotcat1<br>-3.044<br>1.362 | bio14:Plotcat1<br>-2.909<br>3.076 | min_spei12:Plotcat1<br>2.733<br>0.499 | BA:BAcon<br>-2.506<br>0.088 |
| QUEILE | min_spei12:mean_spei12<br>4.279<br>0.156 | BAcon:mean_spei12<br>3.913<br>0.153 | bio12:Plotcat2<br>-3.785<br>0.243 | bio12:min_spei12<br>3.529<br>0.16 | min_spei12:Plotcat1<br>3.337<br>0.141 |
| QUEPYR | bio14:min_spei12<br>-3.424<br>0.291 | BAcon:BA<br>-3.344<br>0.078 | bio14:BA<br>3.324<br>0.101 | min_spei12:BA<br>3.141<br>0.088 | mean_spei12:BA<br>2.646<br>0.398 |
| QUESUB | min_spei12:mean_spei12<br>3.773<br>0.178 | min_spei12:BA<br>3.294<br>0.081 | mean_spei12:Plotcat1<br>3.164<br>1.571 | bio1:Plotcat1<br>2.721<br>0.184 | bio12:min_spei12<br>2.626<br>0.316 |
| ABIALB | BAhetero:mean_spei12<br>4.603<br>0.161 | BA:BAhetero<br>3.2<br>0.048 | BAhetero:min_spei12<br>-2.617<br>0.145 | tmean.djf:bio14<br>-2.511<br>0.064 | BA:mean_spei12<br>-2.505<br>0.123 |
| ACEPSE | bio1:min_spei12<br>-3.595<br>0.171 | bio14:BAcon<br>-3.363<br>0.17 | BAcon:Plotcat1<br>3.267<br>0.68 | mean_spei12:bio1<br>2.869<br>0.323 | bio14:Plotcat1<br>2.499<br>1.068 |
| ALNGLU | BAhetero:min_spei12<br>4.4<br>0.074 | bio1:BAhetero<br>3.881<br>0.106 | bio1:BA<br>-3.328<br>0.095 | Plotcat2:mean_spei12<br>3.176<br>1.731 | bio14:min_spei12<br>2.694<br>0.208 |
| BETPEN | min_spei12:BAcon<br>-6.262<br>0.149 | mean_spei12:BAcon<br>4.923<br>0.212 | bio1:Plotcat2<br>-3.319<br>0.29 | bio1:Plotcat1<br>-2.904<br>0.232 | mean_spei12:Plotcat2<br>2.608<br>0.578 |
| FAGSYL | BA:bio5<br>-4.564<br>0.036 | BAcon:bio5<br>4.126<br>0.038 | bio14:Plotcat2<br>-4.009<br>0.083 | mean_spei12:bio14<br>-3.545<br>0.089 | mean_spei12:BA<br>2.804<br>0.09 |
| FRAEXC | min_spei12:mean_spei12<br>4.415 | min_spei12:Plotcat2<br>3.299 | bio5:BAcon<br>-2.293 | BAhetero:bio5<br>-2.282 | NA<br>NA |

| | | | | | |
|---|---|---|---|---|---|
| | 0.246 | 0.42 | 0.057 | 0.057 | NA |
| PICABI | BA:Plotcat1<br>-8.787<br>0.124 | bio13:mean_spei12<br>6.011<br>0.137 | bio5:bio13<br>-5.751<br>0.055 | mean_spei12:Plotcat1<br>-5.121<br>0.202 | bio13:BA<br>5.111<br>0.08 |
| PINSYL | bio13:mean_spei12<br>-10.705<br>0.108 | min_spei12:BA<br>8.891<br>0.049 | bio13:min_spei12<br>8.876<br>0.091 | mean_spei12:Plotcat1<br>-6.066<br>0.187 | BAcon:min_spei12<br>-5.128<br>0.09 |
| POPNIG | min_spei12:BA<br>2.911<br>0.233 | min_spei12:BAcon<br>-2.712<br>0.318 | bio5:bio12<br>2.671<br>0.184 | BA:Plotcat1<br>-2.616<br>0.418 | NA<br>NA<br>NA |
| POPTRE | bio1:min_spei12<br>-4.326<br>0.124 | mean_spei12:Plotcat1<br>3.93<br>0.524 | bio14:min_spei12<br>3.874<br>0.258 | bio14:mean_spei12<br>-3.32<br>0.322 | mean_spei12:Plotcat2<br>2.352<br>0.962 |
| QUEROB | bio13:BAhetero<br>5.314<br>0.037 | bio13:BA<br>-4.568<br>0.038 | mean_spei12:Plotcat1<br>3.804<br>0.27 | bio5:Plotcat1<br>-3.296<br>0.128 | bio13:min_spei12<br>2.631<br>0.114 |

Table S11d: The five interaction effects with the most important coefficient estimated by the ZTNBMM for each species. Species: Code of the species name. EV: Variable acronym with the importance number indicated. EV t-value: t-value associated with the variable. SE: Standard error associated with the variable.

| Species | EV1<br>EV1 t-value<br>EV 1 SE | EV2<br>EV2 t-value<br>EV 2 SE | EV3<br>EV3 t-value<br>EV 3 SE | EV4<br>EV4 t-value<br>EV 4 SE | EV5<br>EV5 t-value<br>EV 5 SE |
|---|---|---|---|---|---|
| CASSAT | min_spei12:mean_spei12<br>5.089<br>0.186 | bio1:min_spei12<br>3.307<br>0.051 | min_spei12:Plotcat1<br>-2.519<br>0.208 | BA:bio14<br>2.34<br>0.016 | min_spei12:BAhetero<br>2.126<br>0.041 |
| PINHAL | ppet.mean:mean_spei12<br>4.72 | bio5:mean_spei12<br>3.548 | bio5:ppet.mean<br>3.347 | ppet.mean:min_spei12<br>-2.728 | mean_spei12:BAcon<br>2.666 |

| | | | | | |
|---|---|---|---|---|---|
| | 0.218 | 0.21 | 0.111 | 0.055 | 0.125 |
| PINNIG | min_spei12:BAcon<br>-5.117<br>0.06 | mean_spei12:BAcon<br>4.354<br>0.201 | min_spei12:mean_spei12<br>-3.54<br>0.691 | Plotcat2:bio12<br>2.623<br>2.107 | bio5:Plotcat2<br>2.563<br>1.445 |
| PINPINA | bio14:BA<br>-4.587<br>0.024 | Plotcat2:BAhetero<br>-4.298<br>0.032 | min_spei12:BA<br>4.125<br>0.023 | bio1:min_spei12<br>-3.926<br>0.071 | Plotcat2:BA<br>3.357<br>0.037 |
| PINPIN | BAcon:bio5<br>3.415<br>0.047 | Plotcat2:bio5<br>3.099<br>0.741 | Plotcat2:min_spei12<br>2.651<br>0.444 | NA<br>NA<br>NA | NA<br>NA<br>NA |
| QUEILE | mean_spei12:BAcon<br>4.448<br>0.108 | bio1:Plotcat2<br>3.663<br>0.203 | min_spei12:Plotcat2<br>-3.397<br>0.278 | mean_spei12:Plotcat2<br>3.138<br>0.753 | mean_spei12:min_spei12<br>-2.334<br>0.294 |
| QUEPYR | BAcon:bio14<br>-3.33<br>0.058 | Plotcat2:bio1<br>-2.779<br>0.169 | mean_spei12:min_spei12<br>2.682<br>0.979 | BAcon:Plotcat2<br>-2.51<br>0.271 | BAcon:min_spei12<br>2.143<br>0.169 |
| QUESUB | min_spei12:mean_spei12<br>3.069<br>0.191 | min_spei12:Plotcat2<br>2.85<br>0.195 | NA<br>NA<br>NA | NA<br>NA<br>NA | NA<br>NA<br>NA |
| ABIALB | tmean.djf:Plotcat1<br>4.239<br>0.085 | tmean.djf:BAhetero<br>-3.248<br>0.028 | bio14:Plotcat1<br>3.144<br>0.127 | BA:BAhetero<br>-2.181<br>0.026 | tmean.djf:Plotcat2<br>2.181<br>0.083 |
| ACEPSE | Plotcat2:mean_spei12<br>3.793<br>0.373 | min_spei12:Plotcat2<br>-3.179<br>0.432 | BAcon:Plotcat2<br>3.009<br>0.194 | BAcon:bio1<br>2.929<br>0.103 | min_spei12:bio14<br>-2.689<br>0.099 |
| ALNGLU | min_spei12:Plotcat1<br>-3.262<br>0.277 | BA:mean_spei12<br>3.081<br>0.115 | BA:bio1<br>2.904<br>0.047 | min_spei12:BAhetero<br>2.51<br>0.059 | Plotcat1:mean_spei12<br>2.343<br>0.26 |
| BETPEN | Plotcat2:BA<br>3.12 | Plotcat1:mean_spei12<br>-2.541 | Plotcat1:BA<br>2.106 | min_spei12:Plotcat2<br>-2.05 | NA<br>NA |

| | | | | | |
|---|---|---|---|---|---|
| | 0.128 | 0.22 | 0.074 | 0.226 | NA |
| FAGSYL | bio14:min_spei12<br>-4.254<br>0.039 | bio5:min_spei12<br>-2.707<br>0.053 | mean_spei12:Plotcat1<br>2.603<br>0.28 | bio14:Plotcat1<br>-2.012<br>0.124 | NA<br>NA<br>NA |
| FRAEXC | bio12:min_spei12<br>-5.086<br>0.113 | bio12:mean_spei12<br>4.698<br>0.139 | min_spei12:BAhetero<br>3.268<br>0.1 | BAhetero:mean_spei12<br>-3.223<br>0.127 | BAhetero:bio5<br>-2.285<br>0.034 |
| PICABI | BA:BAcon<br>5.18<br>0.027 | bio13:Plotcat2<br>3.7<br>0.134 | Plotcat1:BA<br>-3.348<br>0.043 | bio13:Plotcat1<br>2.799<br>0.123 | NA<br>NA<br>NA |
| PINSYL | BAcon:bio1<br>6.973<br>0.03 | BAcon:mean_spei12<br>5.268<br>0.05 | Plotcat1:mean_spei12<br>4.512<br>0.056 | BA:bio1<br>-3.451<br>0.028 | bio13:Plotcat1<br>2.764<br>0.041 |
| POPTRE | bio1:BA<br>3.143<br>0.075 | BAhetero:BA<br>-2.361<br>0.03 | BA:Plotcat1<br>-2.206<br>0.125 | bio14:mean_spei12<br>-2.161<br>0.102 | NA<br>NA<br>NA |
| QUEPET | bio12:min_spei12<br>-4.624<br>0.04 | bio1:mean_spei12<br>-3.984<br>0.067 | BAcon:bio1<br>3.497<br>0.022 | bio1:Plotcat1<br>-2.936<br>0.048 | bio1:Plotcat2<br>2.638<br>0.086 |
| QUEROB | min_spei12:bio5<br>-3.196<br>0.037 | bio5:Plotcat2<br>2.696<br>0.085 | bio13:BA<br>-2.643<br>0.022 | bio5:Plotcat1<br>-2.622<br>0.079 | bio13:Plotcat1<br>2.446<br>0.067 |

Table S12a: Simple effects significance in binomial model for each species. Species: Code for the species name. Each column represents one variable. When the variable was significant the t-value is reported.

| Species | Census interval | CompetitionINTER | CompetitionINTRA | CompetitionTOTAL | dbh mean (plot) | density | Drought mean | I(Drought mean^2) | (Intercept) | I(Precipitation^2) | I(Temperature^2) | Marginality (LE) | Marginality (TE) | Precipitation | Species growth (plot) | Temperature | Total species growth | Waterbalance |
|---|---|---|---|---|---|---|---|---|---|---|---|---|---|---|---|---|---|---|
| ABIALB | - | -3.49 | - | -6.447 | 13.627 | 5.103 | - | - | -3.447 | -3.241 | - | - | - | - | -9.734 | - | 7.466 | - |
| ACEPSE | - | - | - | -2.316 | 6.869 | 7.165 | -2.565 | - | -5.751 | - | - | - | - | 2.392 | -2.886 | - | - | - |
| ALNGLU | - | - | - | - | 9.591 | 2.577 | - | - | -7.349 | -2.177 | - | -2.53 | 2.047 | - | -6.271 | - | 4.467 | - |
| BETPEN | - | - | -3.543 | -3.644 | 8.319 | 7.349 | -4.369 | - | -4.141 | - | - | - | - | - | -6.3 | 4.839 | 3.749 | - |
| CASSAT | 2.399 | - | - | -10.056 | 21.194 | 13.694 | -10.989 | - | -4.054 | - | -3.799 | - | 3.973 | 5.209 | -10.528 | 3.888 | 10.229 | - |
| FAGSYL | - | - | 26.37 | -12.058 | - | 3.872 | -10.574 | - | -14.291 | 4.749 | - | 2.962 | - | - | -8.507 | - | 3.192 | - |
| FRAEXC | 2 | -6.265 | - | - | 12.397 | 8.091 | - | -2.495 | -4.44 | - | - | 4.229 | - | - | -5.796 | - | 2.486 | - |

| | | | | | | | | | | | | | | | | | | |
|---|---|---|---|---|---|---|---|---|---|---|---|---|---|---|---|---|---|---|
| PICABI | - | - | 12.623 | - | -4.058 | - | - | 2.962 | -8.77 | - | - | 4.22 | 3.735 | -4.022 | -10.222 | -2.958 | 4.238 | - |
| PINHAL | -3.084 | - | 11.139 | -5.065 | 3.01 | 3.416 | 3.477 | - | -7.429 | - | - | -3.041 | - | - | -15.674 | 4.592 | - | 8.116 |
| PINNIG | - | - | 8.926 | -5.075 | - | - | 2.755 | - | - | - | -3.27 | - | - | 3.413 | -11.254 | 3.637 | - | - |
| PINPIN | - | - | 7.43 | -2.856 | 2.069 | - | - | - | - | - | 3.252 | - | - | 4.219 | -8.747 | 3.197 | - | - |
| PINPINA | - | - | - | - | 23.161 | 13.381 | 5.999 | 6.208 | -8.581 | -5.979 | - | 6.421 | -2.709 | - | -15.51 | - | - | - |
| PINSYL | - | - | 7.001 | - | 7.652 | 11.48 | - | - | -10.25 | -2.63 | - | -2.425 | 2.073 | 8.881 | -19.998 | 8.019 | - | - |
| POPNIG | - | - | - | - | 3.466 | 5.066 | - | - | -4.098 | - | - | - | - | - | -3.302 | - | - | - |
| POPTRE | - | - | - | -3.502 | 11.125 | 4.825 | -7.464 | - | -5.993 | - | - | - | - | 3.43 | -5.692 | - | 4.187 | - |
| QUEILE | -3.913 | - | 19.442 | - | -5.4 | - | - | - | -2.314 | - | - | - | - | 5.482 | - | 7.851 | -10.681 | - |
| QUEPYR | - | - | 12.968 | - | - | - | -2.696 | - | -22.406 | - | - | - | - | -4.289 | -8.08 | 4.351 | - | - |

| | | | | | | | | | | | | | | | | | | |
|---|---|---|---|---|---|---|---|---|---|---|---|---|---|---|---|---|---|---|
| QUE ROB | 4.767 | -6.516 | - | -3.288 | 14.73 | 8.433 | -8.427 | - | -5.439 | - | 2.267 | - | - | 3.855 | -15.942 | 2.715 | 13.196 | - |
| QUE SUB | -2.183 | - | 3.884 | - | 4.59 | 3.45 | - | - | -5.306 | - | - | - | - | 2.859 | -8.253 | 2.269 | - | - |

Table S12b. Simple effects significance in zero truncated model for each species. Species: Code for the species name. Each column represents one variable. When the variable was significant the t-value is reported.

| Species | Census interval | CompetitionINTER | CompetitionINTRA | CompetitionTOTAL | dbh mean (plot) | density | Drought mean | I(Drought mean^2) | (Intercept) | I(Precipitation^2) | I(Temperature^2) | I(Waterbalance^2) | Marginality (LE) | Marginality (TE) | Precipitation | Species growth (plot) | Temperature | Total species growth |
|---|---|---|---|---|---|---|---|---|---|---|---|---|---|---|---|---|---|---|
| ABI ALB | - | 3.922 | - | -2.074 | 2.134 | 2.469 | 2.861 | -2.814 | 5.885 | 2.542 | - | - | - | - | -2.068 | - | -2.195 | -11.515 |
| ACE PSE | - | - | -4.503 | 2.958 | - | - | - | - | 4.337 | - | - | - | -4.927 | - | - | 3.4 | - | -4.531 |
| ALN GLU | - | 3.263 | - | -3.207 | 2.297 | -2.736 | - | - | 19.878 | - | - | - | - | -2.214 | - | -7.362 | - | - |
| BET | - | - | -10.473 | - | 7.9 | - | - | - | 13.97 | - | - | - | - | - | - | - | - | - |

| | | | | | | | | | | | | | | | | | | |
|---|---|---|---|---|---|---|---|---|---|---|---|---|---|---|---|---|---|---|
| PEN | | | | | 38 | | | | 1 | | | | 2.663 | | | 7.384 | | |
| CAS SAT | -2.871 | 6.536 | - | -11.518 | 9.201 | 6.06 | 3.893 | -3.232 | 5.522 | - | - | - | - | -2.94 | - | -18.706 | 3.97 | - |
| FAG SYL | -3.809 | - | -7.508 | - | 4.018 | 3.027 | 3.925 | - | 10.725 | - | - | - | - | - | -2.112 | - | -2.206 | -5.695 |
| FRA EXC | 2.839 | 4.361 | -13.558 | - | 8.436 | - | - | - | 32.717 | - | - | - | - | - | -4.957 | -3.956 | - | - |
| PIC ABI | -2.636 | - | - | - | -5.825 | -6.722 | 4.861 | -2.291 | 11.545 | -2.42 | - | - | - | 4.998 | -2.971 | - | - | -16.042 |
| PIN HAL | -8.12 | - | - | - | 4.731 | 7.473 | - | - | 25.362 | - | 3.906 | 2.53 | 2.58 | - | - | - | 2.299 | -17.034 |
| PIN NIG | -3.35 | - | - | - | - | - | -3.947 | - | 8.825 | - | - | - | - | - | - | -2.224 | - | -5.784 |
| PINPIN | -2.793 | - | 5.653 | -3.773 | -4.655 | - | - | - | 7.776 | 2.357 | - | - | 3.2 | - | - | - | - | -9.19 |
| PINPINA | -10.202 | - | - | - | 4.008 | 8.749 | 2.238 | -3.285 | 25.819 | 3.557 | -2.088 | - | -2.13 | - | - | -2.341 | -3.869 | -10.569 |
| PINSYL | -5.742 | - | -10.939 | 3.529 | 5.457 | 9.661 | - | - | 13.979 | -2.945 | -3.825 | - | 2.986 | - | -2.629 | - | - | -14.861 |
| POP | - | 4.453 | - | - | 4.8 | - | - | - | 32.60 | - | - | - | - | - | - | - | - | - |

| | | | | | | | | | | | | | | | | | | |
|---|---|---|---|---|---|---|---|---|---|---|---|---|---|---|---|---|---|---|
| TRE | 2.663 | | | | 79 | | | | 6 | | | | | | | | | 8.301 |
| QUEILE | -4.872 | -2.333 | -2.239 | - | - | 6.538 | -3.051 | - | 9.722 | - | - | - | - | - | - | - | - | -12.383 |
| QUEPET | -2.626 | - | -17.943 | - | 14.768 | 4.952 | - | - | 6.703 | - | - | - | - | - | -5.39 | -5.15 | - | 3.003 |
| QUEPYR | -5.803 | - | 2.743 | - | - | - | 2.806 | -2.943 | 16.028 | - | - | - | - | - | - | - | - | -8.706 |
| QUEROB | -2.062 | 2.222 | - | - | - | 2.08 | - | - | 11.161 | - | 3.259 | - | - | - | -2.98 | 2.264 | - | -9.301 |
| QUESUB | -7.127 | - | -3.839 | - | - | 3.414 | - | -3.674 | 19.853 | 3.537 | 3.859 | - | - | - | - | - | 2.543 | -6.327 |

Table S12c Interaction effects significance in the binomial model for each species. Species: Code for the species name. Each column represents one interaction that includes marginality. LE = Leading edge. TE = Trailing edge. When significant, the t-value is reported.

| Species | Marginality (LE):CompetitionINTRA | Marginality (LE):CompetitionTOTAL | Marginality (LE):Drought mean | Marginality (LE):Precipitation | Marginality (LE):Temperature | Marginality (TE):CompetitionINTRA | Marginality (TE):CompetitionTOTAL | Marginality (TE):Drought mean | Marginality (TE):Precipitation | Marginality (TE):Temperature |
|---|---|---|---|---|---|---|---|---|---|---|
| ABIALB | - | - | 2.327 | - | - | - | - | - | - | - |
| ACEP | - | - | - | - | - | 3.267 | - | - | 2.499 | - |

| | | | | | | | | | | |
|---|---|---|---|---|---|---|---|---|---|---|
| SE | | | | | | | | | | |
| ALN GLU | - | - | 3.176 | - | - | - | - | - | - | - |
| BETP EN | - | - | 2.608 | - | -3.319 | - | - | 2.231 | - | -2.904 |
| CASS AT | - | - | - | - | - | - | - | -2.724 | - | - |
| FAGS YL | - | - | - | -4.009 | - | - | - | - | - | - |
| PICA BI | 2.263 | -4.82 | -2.938 | - | 2.887 | 3.464 | -8.787 | -5.121 | 3.386 | 2.916 |
| PINH AL | - | - | 2.445 | - | - | - | - | - | - | - |
| PINN IG | - | -2.124 | -3.196 | - | 2.056 | - | 3.559 | -2.036 | - | -2.758 |
| PINPI N | - | - | - | - | - | - | - | -2.066 | -2.909 | -3.044 |
| PINPI NA | - | 2.53 | - | - | - | - | - | - | - | 2.232 |
| PINS YL | - | - | 3.024 | - | - | -3.394 | - | -6.066 | -3.968 | - |
| POPN IG | - | - | - | - | - | - | -2.616 | - | - | - |
| POPT RE | - | - | 2.352 | - | - | - | - | 3.93 | - | - |
| QUEI LE | - | - | - | -3.785 | - | - | - | - | -2.843 | - |
| QUEP | - | - | - | - | - | - | - | - | - | -2.403 |

| | | | | | | | | | | |
|---|---|---|---|---|---|---|---|---|---|---|
| YR | | | | | | | | | | |
| QUE ROB | - | - | - | - | - | - | - | 3.804 | - | -3.296 |
| QUES UB | - | - | - | - | - | - | 2.59 | 3.164 | - | 2.721 |

Table S12d Interaction effects significance in the zero truncated model for each species. Species: Code for the species name. Each column represents one interaction that includes marginality. LE = Leading edge. TE = Trailing edge. When significant, the t-value is reported.

| Species | Marginality (LE):CompetitionINTRA | Marginality (LE):CompetitionTOTAL | Marginality (LE):Drought mean | Marginality (LE):Precipitation | Marginality (LE):Temperature | Marginality (TE):CompetitionINTRA | Marginality (TE):CompetitionTOTAL | Marginality (TE):Drought mean | Marginality (TE):Precipitation | Marginality (TE):Temperature |
|---|---|---|---|---|---|---|---|---|---|---|
| ABIA LB | - | - | - | - | 2.181 | - | - | - | 3.144 | 4.239 |
| ACEP SE | 3.009 | - | 3.793 | - | - | -2.386 | - | - | - | - |
| ALN GLU | - | - | - | - | - | - | - | 2.343 | 2.115 | - |
| BETP EN | - | 3.12 | - | - | - | - | 2.106 | -2.541 | - | - |
| FAGS YL | - | - | - | - | - | - | - | 2.603 | -2.012 | - |
| FRAE XC | - | - | - | - | 2.218 | - | - | - | - | - |
| PICA BI | - | - | - | 3.7 | - | - | -3.348 | - | 2.799 | - |
| PINH | - | - | - | - | 2.277 | - | - | - | - | - |

| | | | | | | | | | | |
|---|---|---|---|---|---|---|---|---|---|---|
| AL | | | | | | | | | | |
| PINNIG | - | - | - | 2.623 | 2.563 | - | - | - | - | - |
| PINPIN | - | - | - | - | 3.099 | - | - | - | - | - |
| PINPINA | - | 3.357 | - | - | - | - | - | - | - | - |
| PINSYL | - | - | - | - | - | - | - | 4.512 | 2.764 | - |
| POPTRE | - | - | - | - | - | - | -2.206 | - | - | - |
| QUEILE | - | - | 3.138 | - | 3.663 | - | - | - | - | - |
| QUEPET | - | - | - | - | 2.638 | - | - | - | - | -2.936 |
| QUEPYR | -2.51 | - | - | - | -2.779 | - | - | - | - | - |
| QUEROB | - | - | - | - | 2.696 | - | - | - | 2.446 | -2.622 |
