## Supplemental Figures for "Occurrence but not intensity of mortality rises towards the climatic trailing edge of tree species ranges in European forests"

### Supplementary Information

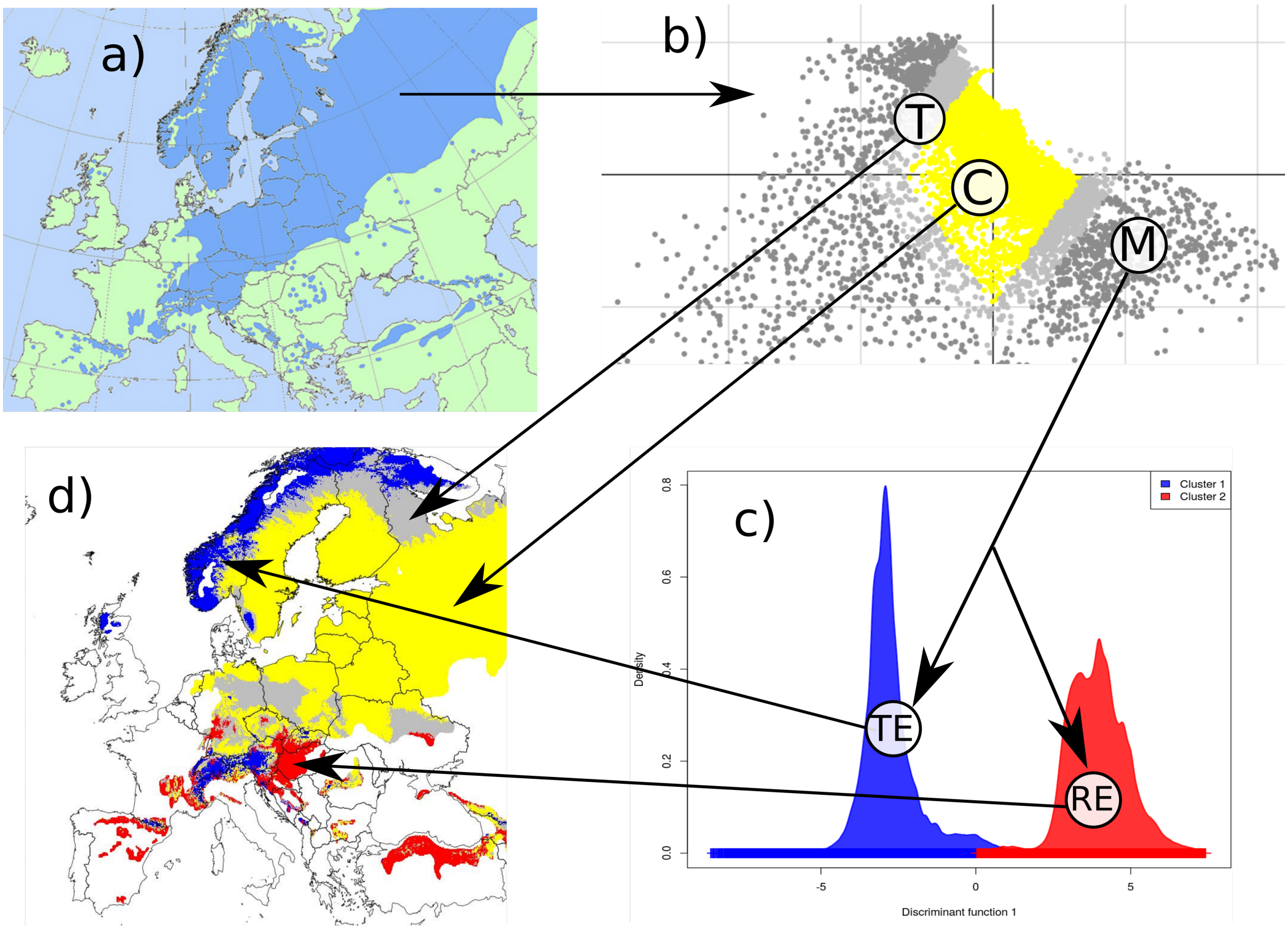


Figure S1: Climatic characterization of the species distribution ranges into core, transition and marginal (leading or trailing) areas. We show an example for *Pinus sylvestris* from EUFORGEN including: a) Species distribution range; b) the climatic variables within the species range are analysed using a Weighted PCA to define three clusters including C = Core areas with C: the lowest weighted scores within the range calculated on the two first axis of the PCA (Core % thresholds (CT) are shown in Table S2 for each species range); M = Marginal areas with M: the largest weighted scores within the range (extremes individuals) calculated on the two first axis of the PCA (margin % thresholds (MT) are shown in Table S2); T = Transition areas with T: weighted scores that fall between the lowest values (core) and largest values (marginal areas); c) results of the Marginal climatic areas that were clustered into: TE trailing edge (TE) and (leading edge (LE) areas (Table S2) with a Discriminant Principal Component Analysis (DPCA).; d) map of the areas defined within the species range with their respective colors. (“Core” in yellow, “trailing edge” in blue, “leading edge” in red and “transition” zone in gray).

Equation S1: Developed form of the functions n1*i,sp* and n2*i,sp* used respectively to model the probability of occurrence of a mortality event in an individual plot *i* during the census interval by species *sp* with a binomial model with a *logit* link in function n1*i,sp* and to model intensity of mortality per year and per hectare in individual plot *i* by species *sp* with a negative binomial model with a *log* link in function n2*i,sp*.$\alpha_{0,sp}$is an intercept term,$\alpha_{country,sp}$is the random country intercept that follows a gaussian distribution (6), $\varepsilon_{i,sp}$is the residual error following a Gaussian distribution (7), $b_{sp}$ is an spatial autocorrelation effect that follows a Matérn distribution (8) with $v_{sp}$ the smoothness and $\rho_{sp}$the shape. (We tested for spatial autocorrelation with the Moran's I statistics ("Moran.I" function of the package "APE" and the function moransI.v of the package "lctools"). Variograms were calculated using the function "correlog" of the package "pgirmess" (Beale, Pleydell, Treglia, & Giraudoux, 2018; Kalogirou, S., & Kalogirou, 2017; Paradis, Claude, & Strimmer, 2004). (Figure S9).

$\beta_{1,sp}$,$\beta_{2,sp}$, $\beta_{3,sp}$, $\beta_{4,sp}$,$\beta_{5,sp}$are the regression coefficients associated with the five covariables, respectively, included in the model (see Table S3). $\beta_{6,sp}$and $\beta_{8,sp}$are the regression coefficients associated with the two climatic variables (see Table S5 and S11) with $\beta_{7,sp}$and$\beta_{9,sp}$the regression coefficients associated with their respective quadratic effect. $\beta_{10,sp}$and $\beta_{12,sp}$are the regression coefficients associated with the two drought-related variables with $\beta_{11,sp}$and$\beta_{13,sp}$ the regression coefficients associated with their respective quadratic effect.$\beta_{14,sp}$and $\beta_{15,sp}$are the regression coefficients associated with the two competition variables effects (Table S3 and S11). $\beta_{16,sp}$are the regression coefficients associated with the climatic marginality effect. $\gamma_1$, $\gamma_{2,sp}$, $\gamma_{3,sp}$, $\gamma_{4,sp}$, $\gamma_{5,sp}$and $\gamma_{6,sp}$are the regression coefficients associated with interaction between the first climatic variable and the second climatic variable, the first drought related variable, the second drought-related variable, the first competition variable, the second competition variable and the climatic marginality, respectively. $\gamma_{7,sp}$, $\gamma_{8,sp}$, $\gamma_{9,sp}$, $\gamma_{10,sp}$and$\gamma_{11,sp}$are the regression coefficients associated with interaction between the second climatic variable and the first drought related variable, the second drought-related variable, the first competition variable, the second competition variable and the climatic marginality, respectively. $\gamma_{12,sp}$, $\gamma_{13,sp}$, $\gamma_{14,sp}$and$\gamma_{15,sp}$are the regression coefficients associated with interaction between the first drought related variable and respectively the second drought-related variable, the first competition variable, the second competition variable and the climatic marginality.$\gamma_{16,sp}$, $\gamma_{17,sp}$and$\gamma_{18,sp}$are the regression coefficients associated with interaction between the second drought related variable and the first competition variable, the second competition variable and the climatic marginality, respectively. $\gamma_{19,sp}$and$\gamma_{20,sp}$are the regression coefficients associated with interaction between the first competition variable and the second competition variable and the climatic marginality, respectively.$\gamma_{21,sp}$is the regression coefficient associated with interaction between the second competition variable and the climatic marginality.

$$
\begin{aligned}
(6)\quad \eta_{i,sp} = {} & \alpha_0 + \alpha_{country,sp} + \varepsilon_{i,sp} + b_{sp} \; + \\
& \beta_{1,sp} \times sqrt(BAIj.plot)_i + \beta_{2,sp} \times sqrt(BAIj.plot.mean)_i + \beta_{3,sp} \times t(dbh.plot.mean)_i + \beta_{4,sp} \times log(treenumber)_i + \\
& \beta_{5,sp} \times log(yearsbetweenstudy)_i + \\
& log(climatic1_i) \times (\beta_{6,sp} + \beta_{7,sp} \times log(climatic1_i) + \gamma_{1,sp} \times log(climatic2_i) + \gamma_{2,sp} \times log(SPEI1_i) + \gamma_{3,sp} \times log(SPEI2_i) + \\
& \quad + \gamma_{4,sp} \times log(competition1_i) + \gamma_{5,sp} \times log(competition2_i) + \gamma_{6,sp} \times log(Marginality_i)) + \\
& log(climatic2_i) \times (\beta_{8,sp} + \beta_{9,sp} \times log(climatic2_i) + \gamma_{7,sp} \times log(SPEI1_i) + \gamma_{8,sp} \times log(SPEI2_i) + \\
& \quad + \gamma_{9,sp} \times log(competition1_i) + \gamma_{10,sp} \times log(competition2_i) + \gamma_{11,sp} \times log(Marginality_i)) + \\
& log(SPEI1_i) \times (\beta_{10,sp} + \beta_{11,sp} \times log(SPEI1_i) + \gamma_{12,sp} \times log(SPEI2_i) + \gamma_{13,sp} \times log(competition1_i) \\
& \quad + \gamma_{14,sp} \times log(competition2_i) + \gamma_{15,sp} \times log(Marginality_i)) + \\
& log(SPEI2_i) \times (\beta_{12,sp} + \beta_{13,sp} \times log(SPEI2_i) + \gamma_{16,sp} \times log(competition1_i) + \gamma_{17,sp} \times log(competition2_i) \\
& \quad + \gamma_{18,sp} \times log(Marginality_i)) + \\
& log(competition1_i) \times (\beta_{14,sp} + \gamma_{19,sp} \times log(competition2_i) + \gamma_{20,sp} \times log(Marginality_i)) + \\
& log(competition2_i) \times (\beta_{15,sp} + \gamma_{21,sp} \times log(Marginality_i)) + log(Marginality_i) \times \beta_{16,sp}
\end{aligned}
$$

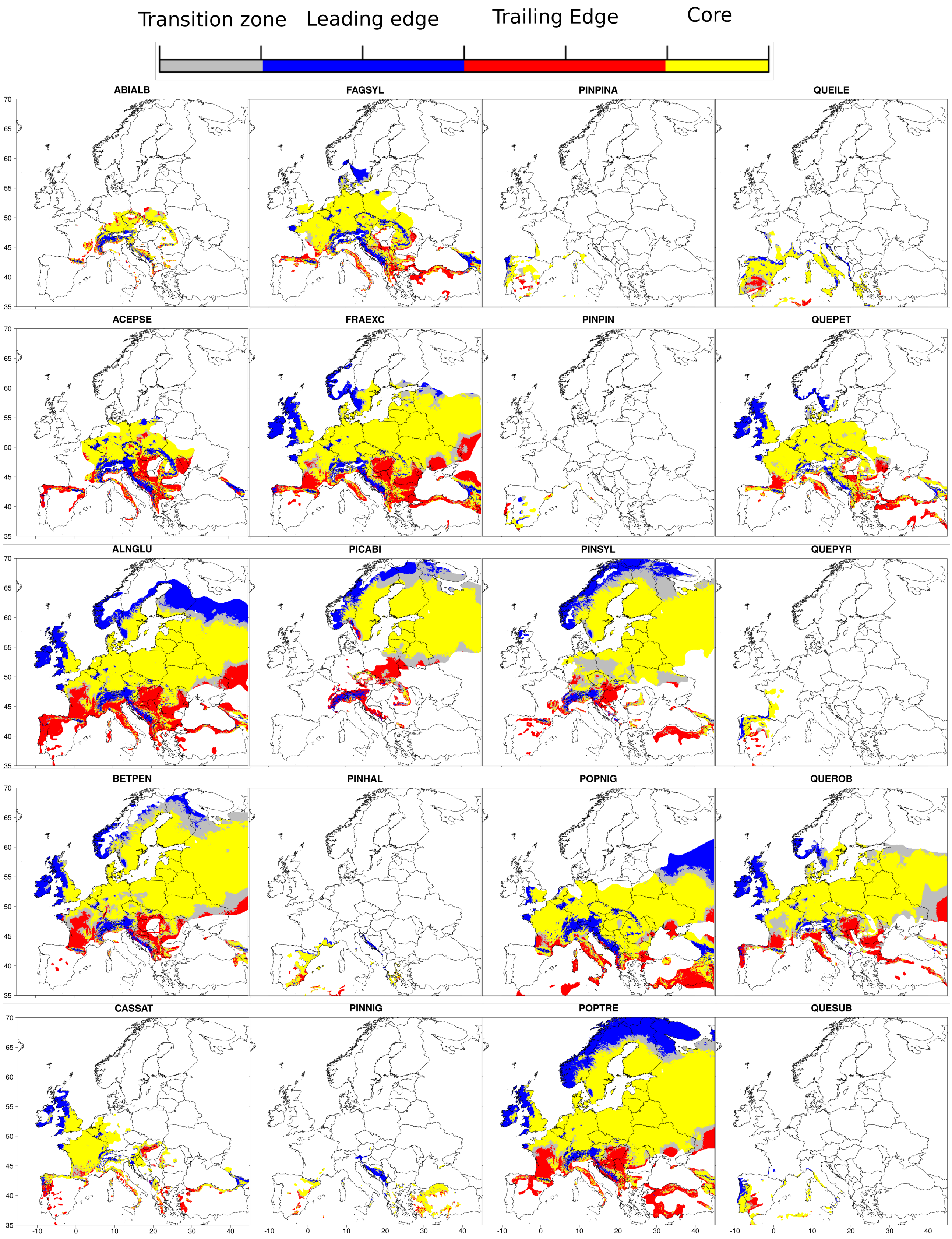


Figure S2: Climatic marginality maps of the 20 tree species studied showing four distribution ranges: core, leading edge, trailing edge and transition areas (in red, blue, red and gray, respectively).

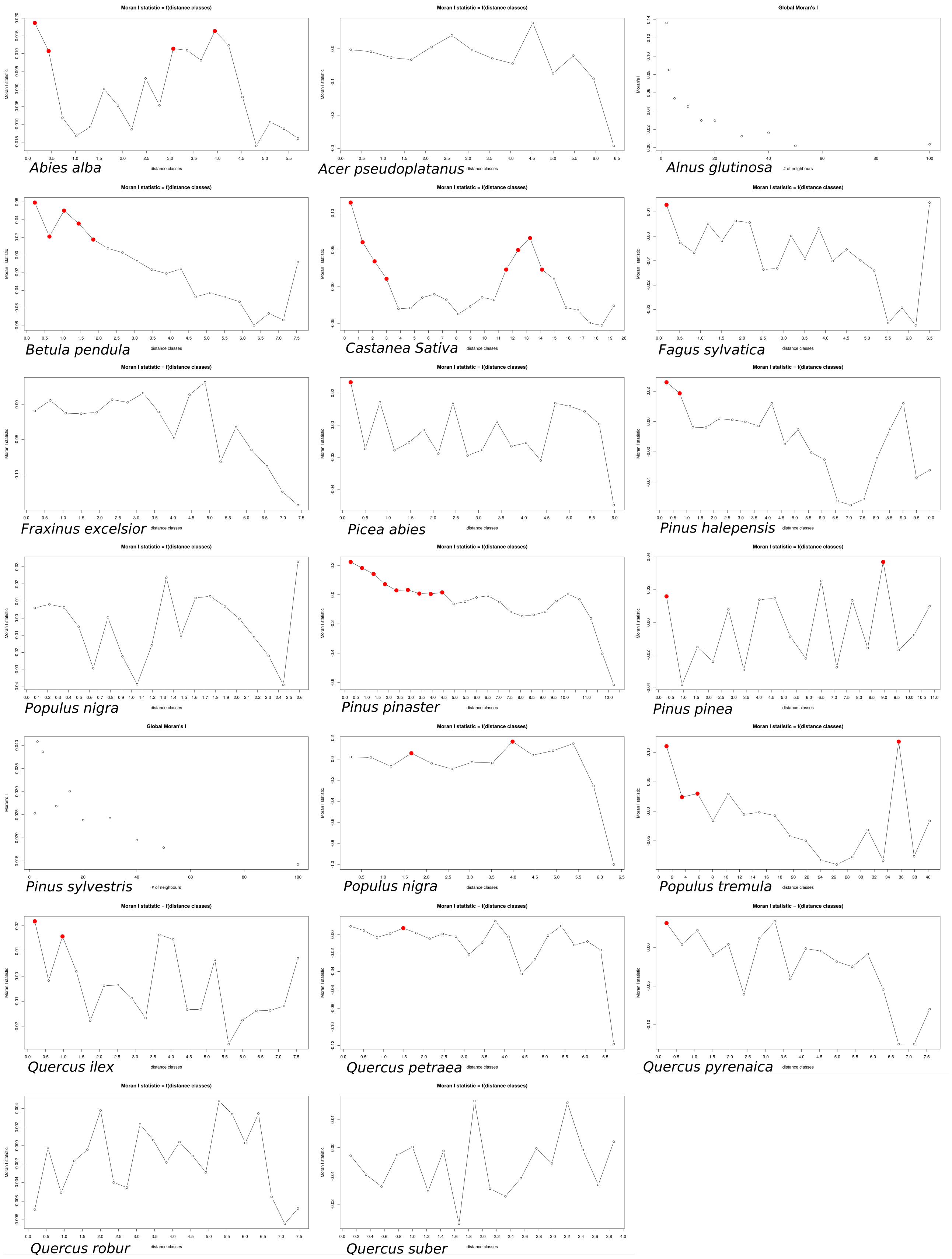


Figure S3: Evaluation of the spatial autocorrelation of observed mortality using variograms and Moran's I statistic from the Pgirmess package (and Lctools when it was not possible). For each species, Moran's I coefficients (Y-axis) are plotted against distance classes (X-axis). Statistically significant values ($p<0.05$) are plotted in red.

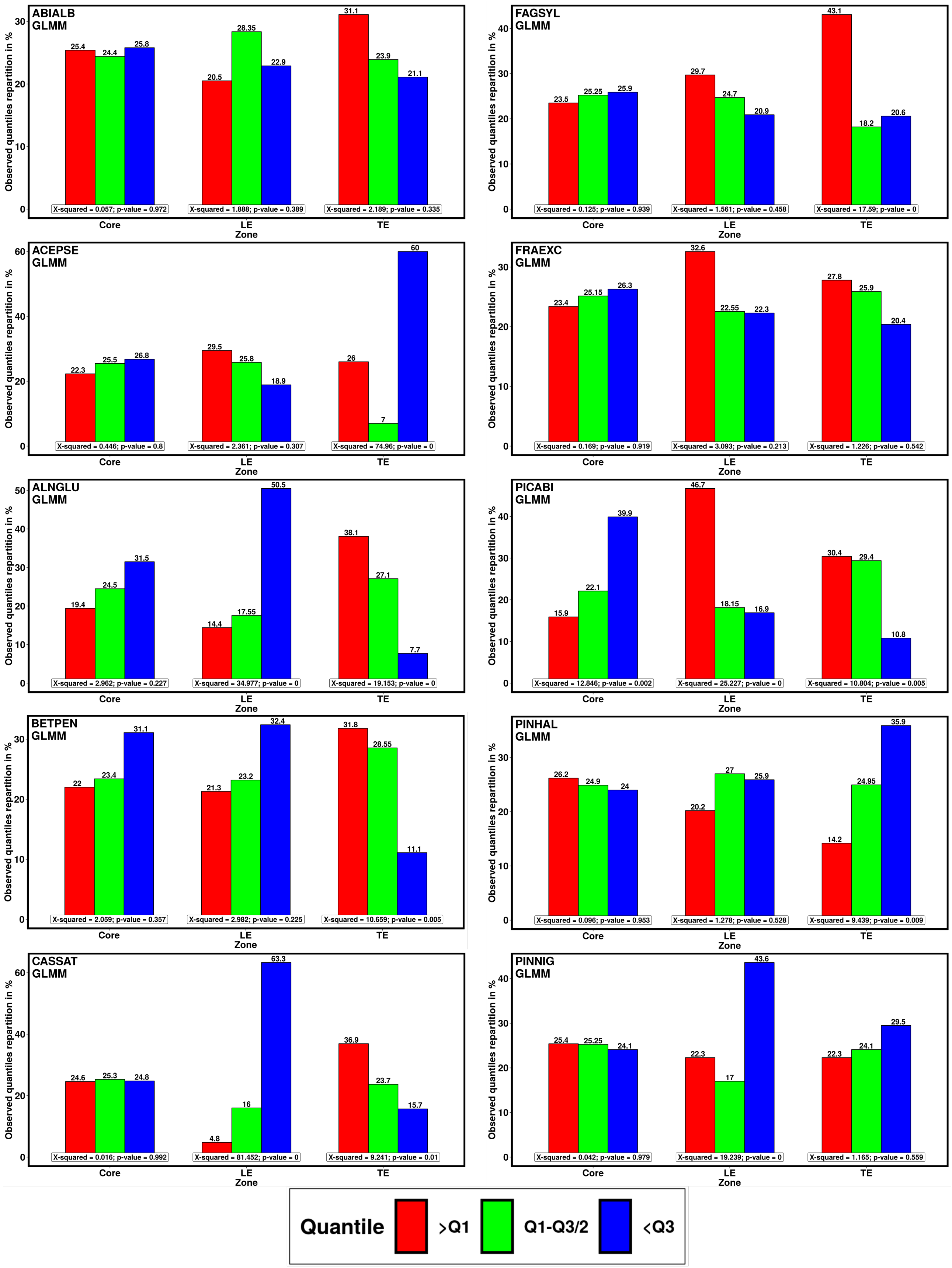

ABIALB GLMM
25.4 24.4 25.8
20.5 28.35 22.9
31.1 23.9 21.1
X-squared = 0.057; p-value = 0.972
X-squared = 1.888; p-value = 0.389
X-squared = 2.189; p-value = 0.335
FAGSYL GLMM
23.5 25.25 25.9
29.7 24.7 20.9
43.1 18.2 20.6
X-squared = 0.125; p-value = 0.939
X-squared = 1.561; p-value = 0.458
X-squared = 17.59; p-value = 0
ACEPSE GLMM
22.3 25.5 26.8
29.5 25.8 18.9
26 7 60
X-squared = 0.446; p-value = 0.8
X-squared = 2.361; p-value = 0.307
X-squared = 74.96; p-value = 0
FRAEXC GLMM
23.4 25.15 26.3
32.6 22.55 22.3
27.8 25.9 20.4
X-squared = 0.169; p-value = 0.919
X-squared = 3.093; p-value = 0.213
X-squared = 1.226; p-value = 0.542
ALNGLU GLMM
19.4 24.5 31.5
14.4 17.55 50.5
38.1 27.1 7.7
X-squared = 2.962; p-value = 0.227
X-squared = 34.977; p-value = 0
X-squared = 19.153; p-value = 0
PICABI GLMM
15.9 22.1 39.9
46.7 18.15 16.9
30.4 29.4 10.8
X-squared = 12.846; p-value = 0.002
X-squared = 25.227; p-value = 0
X-squared = 10.804; p-value = 0.005
BETPEN GLMM
22 23.4 31.1
21.3 23.2 32.4
31.8 28.55 11.1
X-squared = 2.059; p-value = 0.357
X-squared = 2.982; p-value = 0.225
X-squared = 10.659; p-value = 0.005
PINHAL GLMM
26.2 24.9 24
20.2 27 25.9
14.2 24.95 35.9
X-squared = 0.096; p-value = 0.953
X-squared = 1.278; p-value = 0.528
X-squared = 9.439; p-value = 0.009
CASSAT GLMM
24.6 25.3 24.8
4.8 16 63.3
36.9 23.7 15.7
X-squared = 0.016; p-value = 0.992
X-squared = 81.452; p-value = 0
X-squared = 9.241; p-value = 0.01
PINNIG GLMM
25.4 25.25 24.1
22.3 17 43.6
22.3 24.1 29.5
X-squared = 0.042; p-value = 0.979
X-squared = 19.239; p-value = 0
X-squared = 1.165; p-value = 0.559
Observed quantiles repartition in %
Core
LE
TE
Zone
Quantile
>Q1
Q1-Q3/2
<Q3

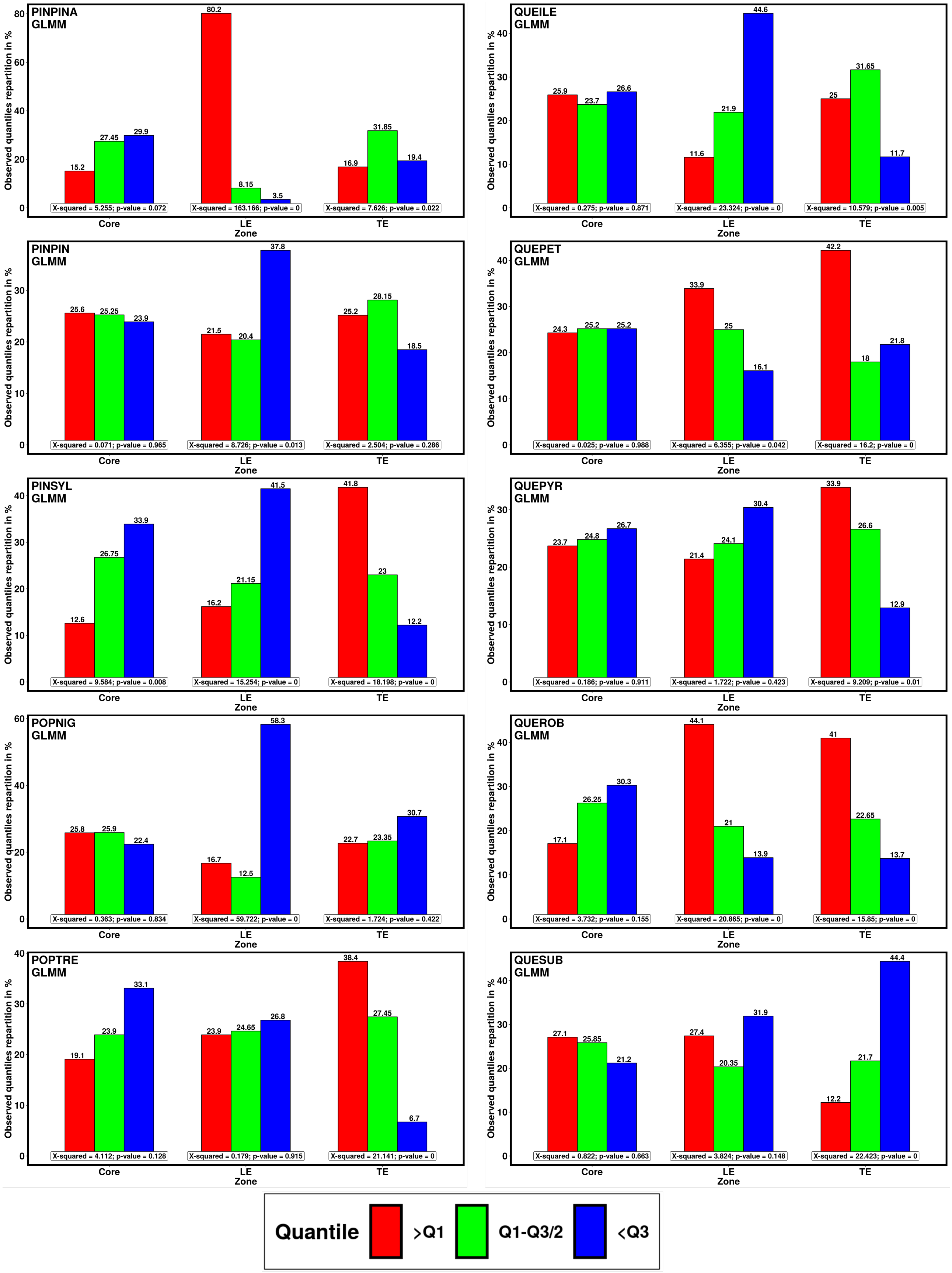

PINPINA
GLMM
80.2
15.2
27.45
29.9
8.15
3.5
16.9
31.85
19.4
X-squared = 5.255; p-value = 0.072
X-squared = 163.166; p-value = 0
X-squared = 7.626; p-value = 0.022
QUEILE
GLMM
25.9
23.7
26.6
11.6
21.9
44.6
25
31.65
11.7
X-squared = 0.275; p-value = 0.871
X-squared = 23.324; p-value = 0
X-squared = 10.579; p-value = 0.005
PINPIN
GLMM
25.6
25.25
23.9
21.5
20.4
37.8
25.2
28.15
18.5
X-squared = 0.071; p-value = 0.965
X-squared = 8.726; p-value = 0.013
X-squared = 2.504; p-value = 0.286
QUEPET
GLMM
24.3
25.2
25.2
33.9
25
16.1
42.2
18
21.8
X-squared = 0.025; p-value = 0.988
X-squared = 6.355; p-value = 0.042
X-squared = 16.2; p-value = 0
PINSYL
GLMM
12.6
26.75
33.9
16.2
21.15
41.5
41.8
23
12.2
X-squared = 9.584; p-value = 0.008
X-squared = 15.254; p-value = 0
X-squared = 18.198; p-value = 0
QUEPYR
GLMM
23.7
24.8
26.7
21.4
24.1
30.4
33.9
26.6
12.9
X-squared = 0.186; p-value = 0.911
X-squared = 1.722; p-value = 0.423
X-squared = 9.209; p-value = 0.01
POPNIG
GLMM
25.8
25.9
22.4
16.7
12.5
58.3
22.7
23.35
30.7
X-squared = 0.363; p-value = 0.834
X-squared = 59.722; p-value = 0
X-squared = 1.724; p-value = 0.422
QUEROB
GLMM
17.1
26.25
30.3
44.1
21
13.9
41
22.65
13.7
X-squared = 3.732; p-value = 0.155
X-squared = 20.865; p-value = 0
X-squared = 15.85; p-value = 0
POPTRE
GLMM
19.1
23.9
33.1
23.9
24.65
26.8
38.4
27.45
6.7
X-squared = 4.112; p-value = 0.128
X-squared = 0.179; p-value = 0.915
X-squared = 21.141; p-value = 0
QUESUB
GLMM
27.1
25.85
21.2
27.4
20.35
31.9
12.2
21.7
44.4
X-squared = 0.822; p-value = 0.663
X-squared = 3.824; p-value = 0.148
X-squared = 22.423; p-value = 0
Observed quantiles repartition in %
Core
LE
TE
Zone
Quantile
>Q1
Q1-Q3/2
<Q3

Fig S4a: Quantile repartition of the probability of mortality occurrence predictions (Fig 4) between lowest values (blue bars), medium values (green bars) and highest values (red bars) according to plots location (core, leading edge and trailing edge) and for each species. This distribution was used to test for heterogeneity of the distribution among groups with a χ-square test. Under the assumption of no spatial structure in mortality occurrence probability (null hypothesis), we expect the distribution to be evenly distributed as follow : For each area (core, leading edge, trailing edge) we expect the first quantile (largest probability, red bars) to represent 25% of the values, the fourth quartile (lowest probability, blue bars) to represent 25% of the values and the second and third quartile (intermediate probability, green bars) to represent 50% of the values. Note that for the sake of readability, we represented the green bars values divided by two. P-values $< 0.05$ indicate that predicted mortality are different than expected. χ-square statistics and p-values are written at the bottom of each bar groups.

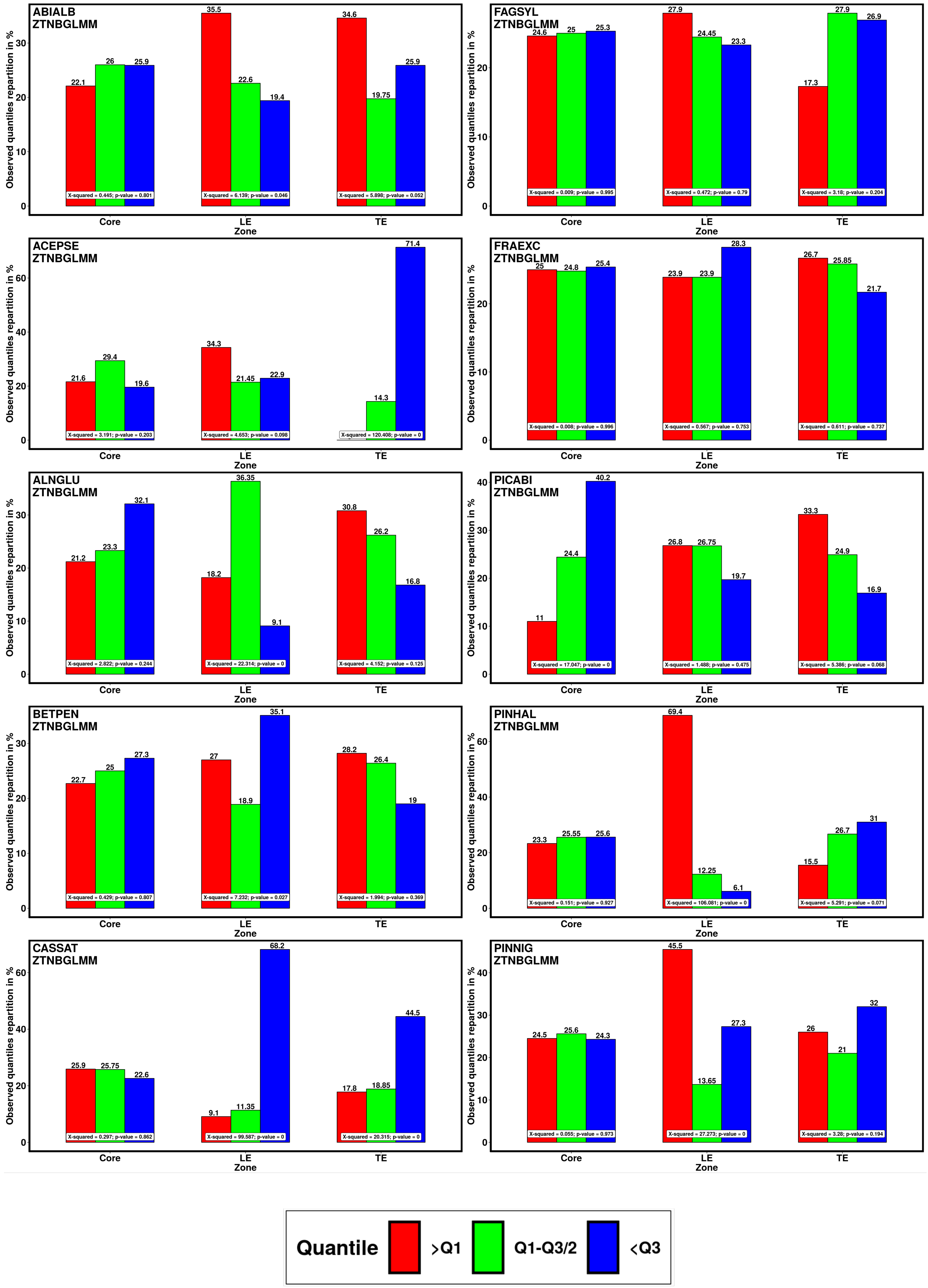

ABIALB ZTNBGLMM
22.1 26 25.9 35.5 22.6 19.4 34.6 19.75 25.9
X-squared = 0.445; p-value = 0.801
X-squared = 6.139; p-value = 0.046
X-squared = 5.898; p-value = 0.052
FAGSYL ZTNBGLMM
24.6 25 25.3 27.9 24.45 23.3 17.3 27.9 26.9
X-squared = 0.009; p-value = 0.995
X-squared = 0.472; p-value = 0.79
X-squared = 3.18; p-value = 0.204
ACEPSE ZTNBGLMM
21.6 29.4 19.6 34.3 21.45 22.9 14.3 71.4
X-squared = 3.191; p-value = 0.203
X-squared = 4.653; p-value = 0.098
X-squared = 120.408; p-value = 0
FRAEXC ZTNBGLMM
25 24.8 25.4 23.9 23.9 28.3 26.7 25.85 21.7
X-squared = 0.008; p-value = 0.996
X-squared = 0.567; p-value = 0.753
X-squared = 0.611; p-value = 0.737
ALNGLU ZTNBGLMM
21.2 23.3 32.1 18.2 36.35 9.1 30.8 26.2 16.8
X-squared = 2.822; p-value = 0.244
X-squared = 22.314; p-value = 0
X-squared = 4.152; p-value = 0.125
PICABI ZTNBGLMM
11 24.4 40.2 26.8 26.75 19.7 33.3 24.9 16.9
X-squared = 17.047; p-value = 0
X-squared = 1.488; p-value = 0.475
X-squared = 5.386; p-value = 0.068
BETPEN ZTNBGLMM
22.7 25 27.3 27 18.9 35.1 28.2 26.4 19
X-squared = 0.429; p-value = 0.807
X-squared = 7.232; p-value = 0.027
X-squared = 1.994; p-value = 0.369
PINHAL ZTNBGLMM
23.3 25.55 25.6 69.4 12.25 6.1 15.5 26.7 31
X-squared = 0.151; p-value = 0.927
X-squared = 106.081; p-value = 0
X-squared = 5.291; p-value = 0.071
CASSAT ZTNBGLMM
25.9 25.75 22.6 9.1 11.35 68.2 17.8 18.85 44.5
X-squared = 0.297; p-value = 0.862
X-squared = 99.587; p-value = 0
X-squared = 20.315; p-value = 0
PINNIG ZTNBGLMM
24.5 25.6 24.3 45.5 13.65 27.3 26 21 32
X-squared = 0.055; p-value = 0.973
X-squared = 27.273; p-value = 0
X-squared = 3.28; p-value = 0.194
Observed quantiles repartition in %
Core
LE
TE
Zone
Quantile
>Q1
Q1-Q3/2
<Q3

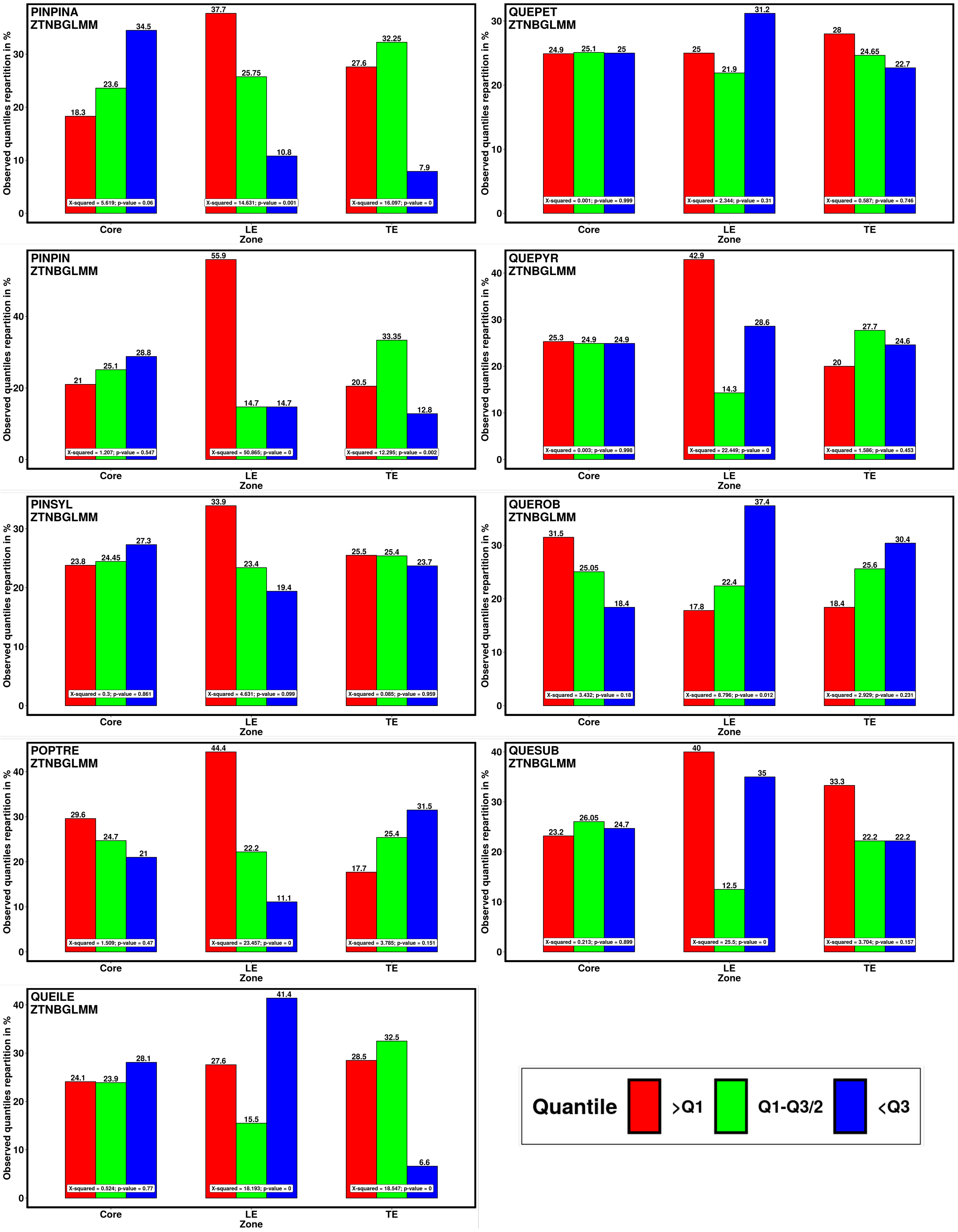
PINPINA
ZTNBGLMM
QUEPET
ZTNBGLMM
PINPIN
ZTNBGLMM
QUEPYR
ZTNBGLMM
PINSYL
ZTNBGLMM
QUEROB
ZTNBGLMM
POPTRE
ZTNBGLMM
QUESUB
ZTNBGLMM
QUEILE
ZTNBGLMM
Observed quantiles repartition in %
Core
LE
TE
Zone
X-squared = 5.619; p-value = 0.06
X-squared = 14.631; p-value = 0.001
X-squared = 16.097; p-value = 0
X-squared = 0.001; p-value = 0.999
X-squared = 2.344; p-value = 0.31
X-squared = 0.587; p-value = 0.746
X-squared = 1.207; p-value = 0.547
X-squared = 50.865; p-value = 0
X-squared = 12.295; p-value = 0.002
X-squared = 0.003; p-value = 0.998
X-squared = 22.449; p-value = 0
X-squared = 1.586; p-value = 0.453
X-squared = 0.3; p-value = 0.861
X-squared = 4.631; p-value = 0.099
X-squared = 0.085; p-value = 0.959
X-squared = 3.432; p-value = 0.18
X-squared = 8.796; p-value = 0.012
X-squared = 2.929; p-value = 0.231
X-squared = 1.509; p-value = 0.47
X-squared = 23.457; p-value = 0
X-squared = 3.785; p-value = 0.151
X-squared = 0.213; p-value = 0.899
X-squared = 25.5; p-value = 0
X-squared = 3.704; p-value = 0.157
X-squared = 0.524; p-value = 0.77
X-squared = 18.193; p-value = 0
X-squared = 18.547; p-value = 0
Quantile
>Q1
Q1-Q3/2
<Q3

Fig S4b: Quantile repartition of the annual intensity of mortality predictions (Fig 4) between lowest values (blue bars), medium values (green bars) and highest values (red bars) according to plots location (core, leading edge and trailing edge) and for each species. This distribution was used to test for heterogeneity of the distribution among groups with a $\chi$-square test. Under the assumption of no spatial structure in mortality occurrence probability (null hypothesis), we expect the distribution to be evenly distributed as follow : For each area (core, leading edge, trailing edge) we expect the first quantile (largest probability, red bars) to represent 25% of the values, the fourth quartile (lowest probability, blue bars) to represent 25% of the values and the second and third quartile (intermediate probability, green bars) to represent 50% of the values. Note that for the sake of readability, we represented the green bars values divided by two. P-values $< 0.05$ indicate that predicted mortality are different than expected. $\chi$-square statistics and p-values are written at the bottom of each bar groups.

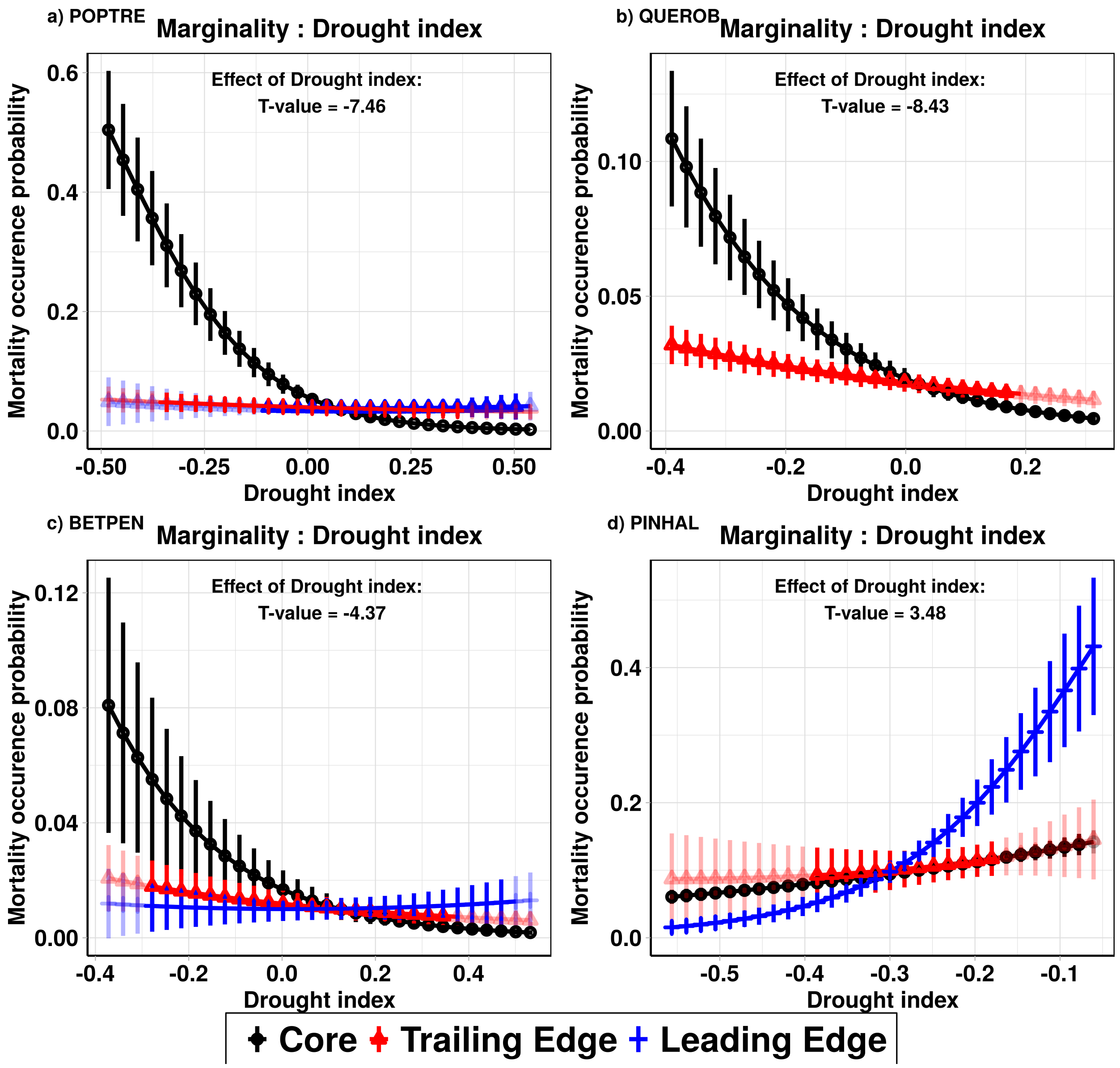


Figure S5: Statistical interactions between SPEI index (X-axis) and mortality occurrence (expressed as probability, Y-axis) across the core, trailing and leading climatic margins defined for three temperate species a) *Populus tremula*, b) *Quercus robur* and c) *Betula pendula* and for the Mediterranean species d)*Pinus halepensis*. Black, red and blue lines represent populations at the core, trailing and leading edge the distribution range, respectively. Predictions within the ranges of the environmental gradients covered by the species are shown by solid colors and extrapolations outside the environmental gradients covered by the species are shown in light colors.

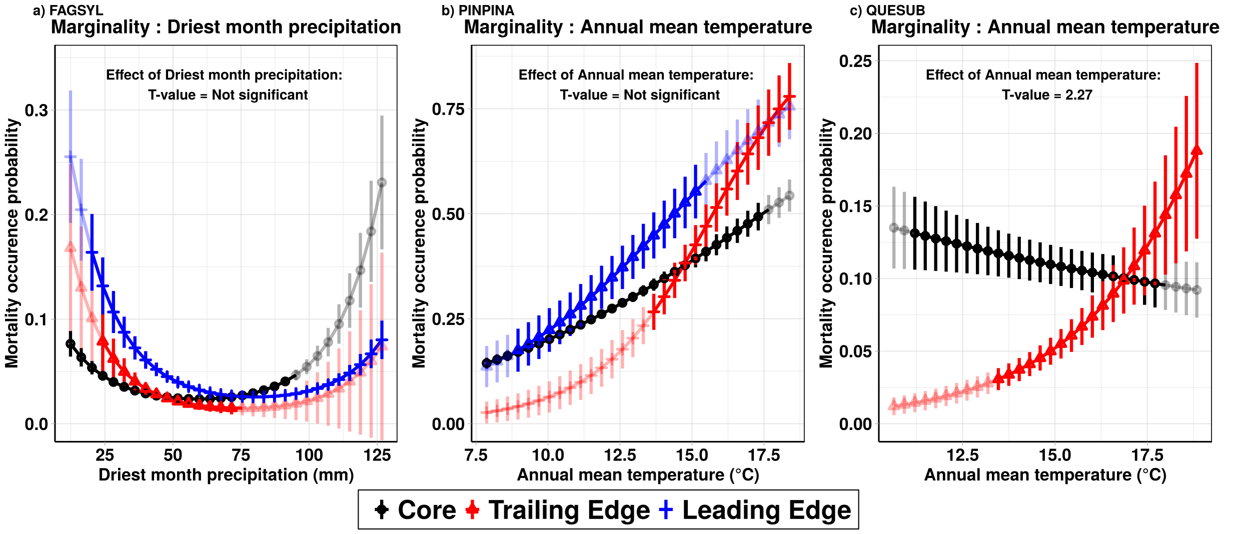


Figure S6: Model interactions between climatic variables and occurrence of mortality (expressed as probability, Y-axis) across the core, trailing and leading climatic margins defined for a) *Fagus sylvatica*, b) *Pinus pinaster* and c), *Quercus suber*. Black, red and blue lines represent populations at the core, trailing and leading edge of the distribution range, respectively. Predictions within the ranges of the environmental gradients covered by the species are shown by solid colors and extrapolations outside the environmental gradients covered by the species are shown in light colors.

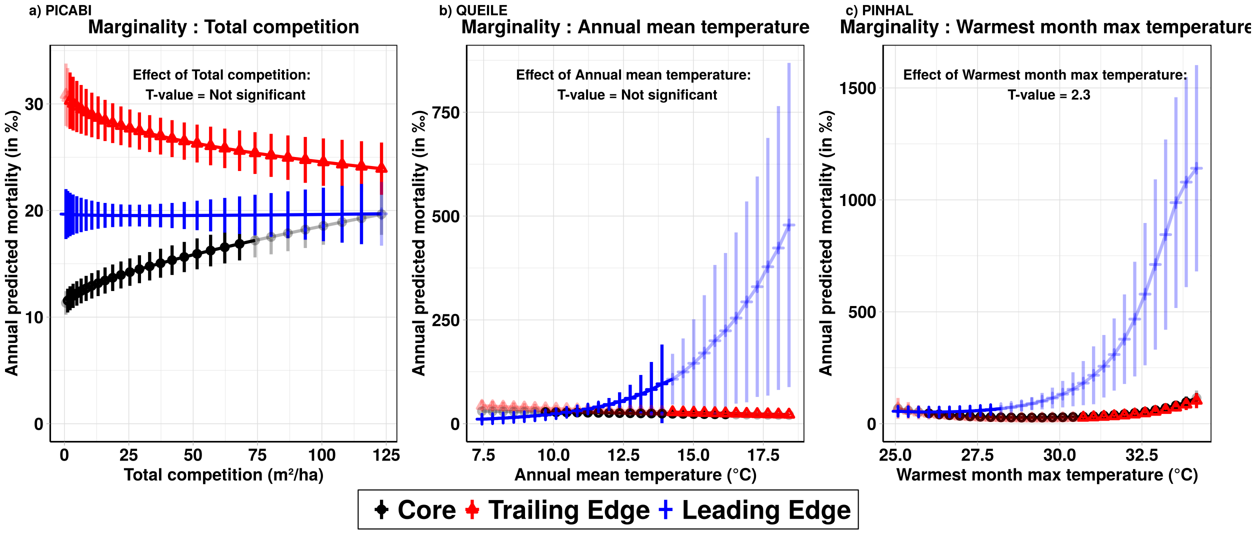


Figure S7: Statistical interactions between several climatic and competition variables and the intensity of predicted mortality by year and by plot (‰) across the core, trailing and leading edge of a) *Picea abies*, b) *Quercus ilex* and c) *Pinus halepensis*. Black, red and blue lines represent populations at the core, trailing and leading edge the distribution range, respectively. Predictions within the ranges of the environmental gradients covered by the species are shown by solid colors and extrapolations outside the environmental gradients covered by the species are shown in light colors.

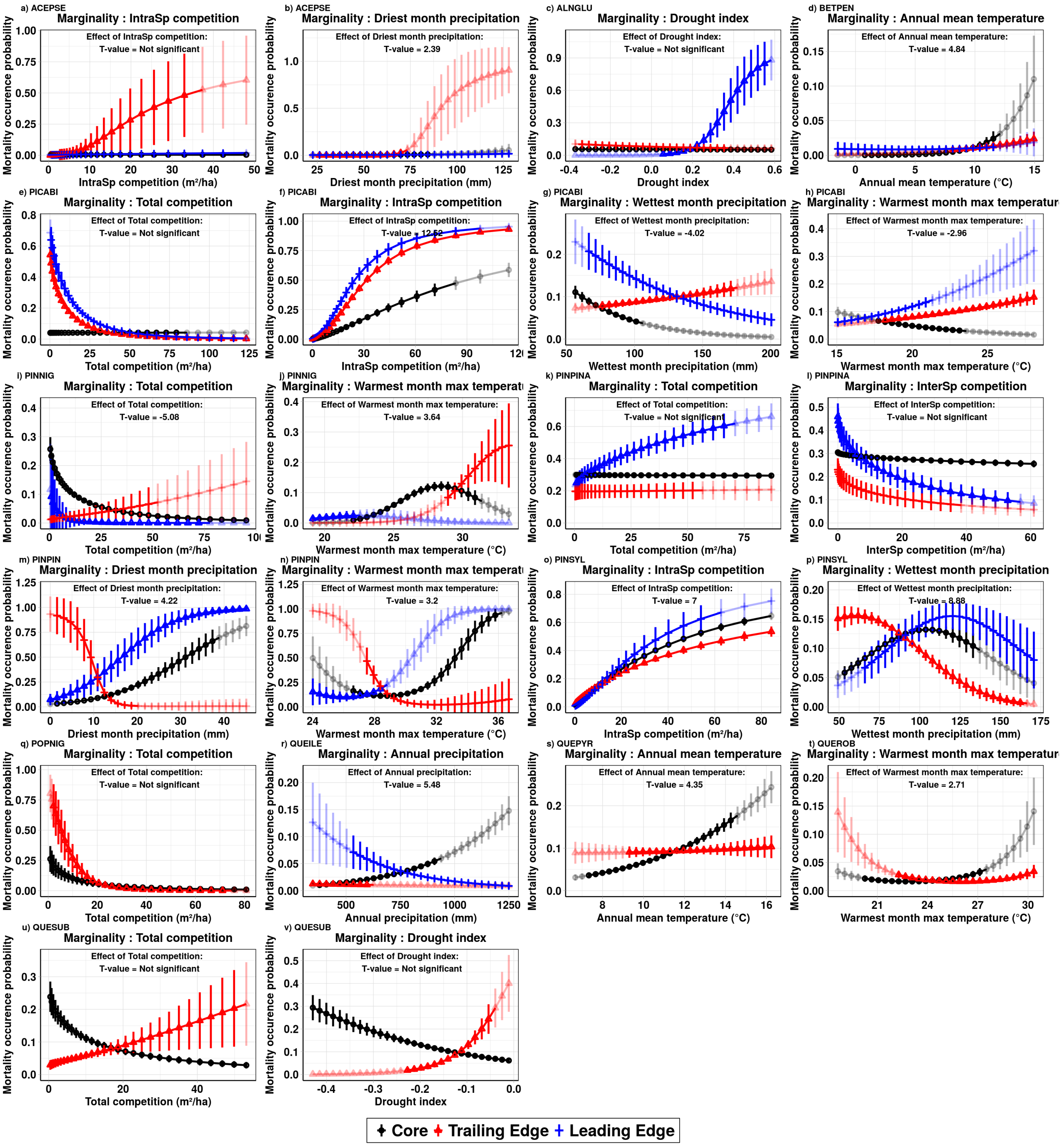


Figure S8: Other significant interactions between several climatic variables (X-axis) and mortality occurrence of mortality (expressed as probability, Y-axis) across the core, trailing and leading edge for all species. Black, red and blue lines represent populations at the core, trailing and leading edge the distribution range, respectively. Predictions within the ranges of the environmental gradients covered by the species are shown by solid colors and extrapolations outside the environmental gradients covered by the species are shown in light colors.

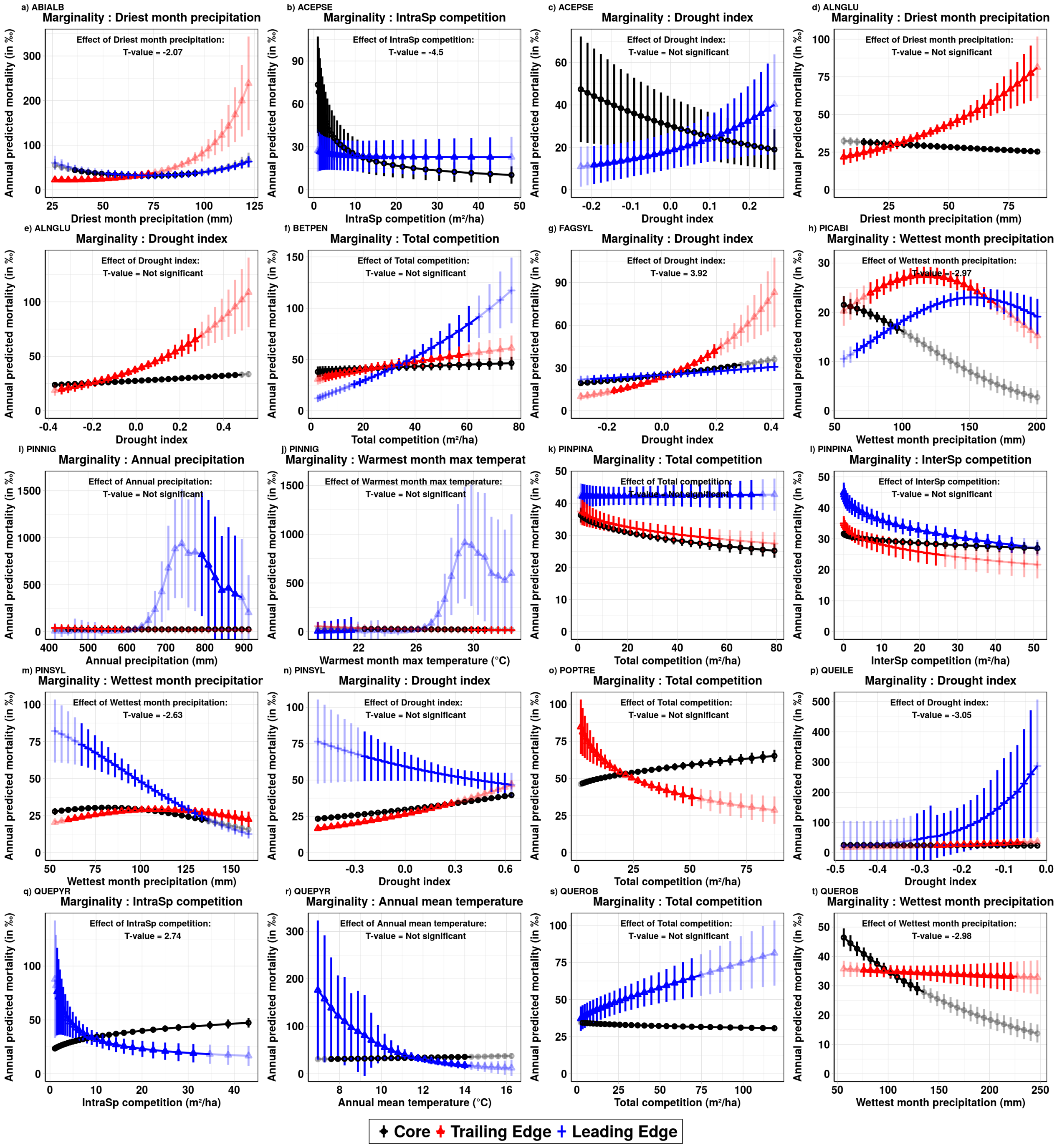


Figure S9: Other significant interactions between several climatic variables (X-axis) and the intensity of predicted mortality (expressed as proportion (‰), by year and by plot, Y-axis) across the core, trailing and leading edge for all species. Black, red and blue lines represent populations at the core, trailing and leading edge the distribution range, respectively. Predictions within the ranges of the environmental gradients covered by the species are shown by solid colors and extrapolations outside the environmental gradients covered by the species are shown in light colors.
